# SHARK enables homology assessment in unalignable and disordered sequences

**DOI:** 10.1101/2023.06.26.546490

**Authors:** Chi Fung Willis Chow, Soumyadeep Ghosh, Anna Hadarovich, Agnes Toth-Petroczy

**Affiliations:** Max Planck Institute of Molecular Cell Biology and Genetics, Pfotenhauerstrasse 108, 01307 Dresden, Germany; Center for Systems Biology Dresden, Pfotenhauerstrasse 108, 01307 Dresden, Germany; Cluster of Excellence Physics of Life, TU Dresden, 01062 Dresden, Germany

**Keywords:** intrinsically disordered protein regions, homology detection, sequence to function paradigm, machine learning

## Abstract

Intrinsically disordered regions (IDRs) are structurally flexible protein segments with regulatory functions in multiple contexts, such as in the assembly of biomolecular condensates.

Since IDRs undergo more rapid evolution than ordered regions, identifying homology of such poorly conserved regions remains challenging for state-of-the-art alignment-based methods that rely on position-specific conservation of residues. Thus, systematic functional annotation and evolutionary analysis of IDRs have been limited, despite comprising ∼21% of proteins.

To accurately assess homology between unalignable sequences, we developed an alignment-free sequence comparison algorithm, SHARK (Similarity/Homology Assessment by Relating K-mers). We trained SHARK-dive, a machine learning homology classifier, which achieved superior performance to standard alignment in assessing homology in unalignable sequences, and correctly identified dissimilar IDRs capable of functional rescue in IDR-replacement experiments reported in the literature.

SHARK-dive not only predicts functionally similar IDRs, but also identifies cryptic sequence properties and motifs that drive remote homology, thereby facilitating systematic analysis and functional annotation of the unalignable protein universe.

## Introduction

A key goal in biology is to decipher the function of all proteins. Understanding of sequence-function relationships by identifying sequences/characteristics conferring a particular function would ultimately facilitate rational protein design^1^. Whilst it is impossible to experimentally annotate the function of every sequence due to the size of the protein universe (the 2022_03 release of UniProt has >227 million proteins)^2^, homology-based protein function inference has aided in this effort^3–5^. Specifically, sequence alignment algorithms such as Smith-Waterman local alignment (SW)^6^, from which BLAST^7,8^ was developed as a faster heuristic, and hidden Markov model algorithms such as HMMer^9^ achieved great success in identifying homologous sequences. To date, most conserved regions have been grouped into domain families and catalogued in databases such as Pfam, where the function of the member sequences are predicted to be similar^10,11^. Nevertheless, a significant fraction of residues cannot be easily aligned to domain families (henceforth referred to as unalignable sequences). Recent Pfam releases have indicated a plateau in the percentage of UniProt sequences (∼73%) and residues (∼53%) mapped to domains^10^, suggesting that a large proportion of sequences and residues is and will remain challenging for alignment-based analyses.

Alignment-based algorithms are also unsuitable for most intrinsically disordered regions (IDRs). These are flexible sequences that evolve more rapidly than ordered domains since they lack constraints for a stable 3D structure^12–15^. As a result, they harbor more substitutions and insertions/deletions (InDels) between homologs than ordered sequences^16,17^. As illustrated with the multiple sequence alignment of *S. cerevisiae* Ded1p orthologs (Fig. 1a): the N- and C-terminal IDRs of Ded1p are poorly aligned, indicated by the large fraction of gaps and amino acid mismatches. Additionally, IDRs often contain biased amino acid compositions and repetitive regions which are not amenable to alignment-based algorithms^18,19^. Consequently, IDRs are largely ignored when identifying homologous sequences, where sensitivity is mostly derived from alignment of conserved domains.

**Figure 1.**
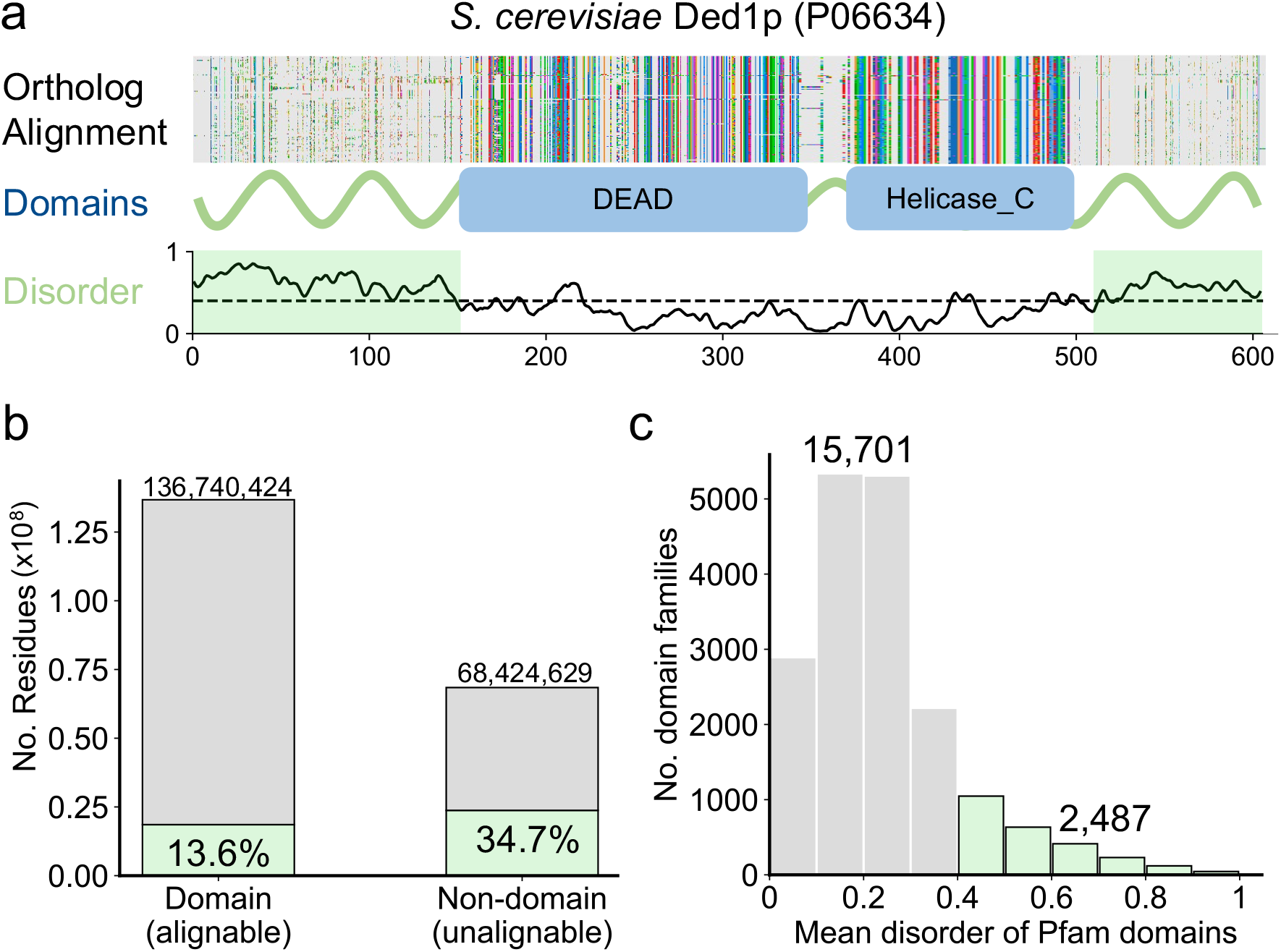
IDRs are common in the proteome and difficult to align. **a**. Multiple sequence alignment of Ded1p orthologs. The disordered N- and C-terminals (shaded in green) contain large number of gaps (grey) in the alignment and a lack of domain annotation due to poor alignment. On the contrary, Pfam domains (blue boxes) correspond to well-aligned, ordered regions with conserved sequences. **b**. Of all ∼205 million residues in Swiss-Prot, 33% do not belong to Pfam domains, representing unalignable residues without systematic functional annotation. Disordered residues are enriched in non-domain (unalignable) regions with greater than two-fold relative enrichment compared to domains. **c**. Disorder is poorly represented in Pfam database (release 34.0). Of the seed sequences used to construct each domain-specific profile HMM of 18188 domain families analyzed, only 2487 families (14%) contain seed sequences with at least 0.4 mean disorder (see Methods).

Whereas IDRs have previously been thought of simply as flexible linkers between the functional structured domains, rapidly increasing evidence has demonstrated the importance of IDRs in modulating protein function^15,20^. Due to their inherent flexibility, multi-valency and ability to sample multiple conformations, they mediate a wide variety of functions including molecular recognition, protein modification, molecular assembly^15^ and biomolecular condensate formation^21–26^. For example, the IDRs of Ded1p play an integral role in regulating its condensation and translation repression^27^. Despite ample evidence of IDRs as functional regions under conservation, homology assessment of IDR remains a bottleneck.

With the recent advances in deep learning and natural language processing, methods such as DeepBLAST^28^, ProtENN^29^ and DEDAL^30^ revisited the pairwise alignment algorithms developed in the 1980-90s. These models harnessed large amounts of sequence data and computing power to improve on alignment accuracy by optimizing parameters such as substitution scores and gap penalties. Whilst they showed improvements in detection and alignment of homologous sequence pairs with low sequence identity, they assume that an accurate alignment exists, and assessed performance primarily on ordered domains^28–30^.

The underlying principles of alignment require a collinear correspondence between residues and assumes that the two sequences can be connected via a mutational path of substitutions, insertions and deletions^31^. However, beyond a certain level of divergence, this approach becomes inaccurate. In rapidly evolving IDRs, only short motifs (SLiMs)^32–34^ dispersed along the sequence or amino acid composition are conserved across orthologs. Notably, these conserved features nested in disordered regions can be essential for protein function^35,36^, and would often be overlooked using alignment methods.

Here we developed SHARK (Similarity/Homology Assessment by Relating K-mers), an alignment-free algorithm that compares the overall amino acid composition and short motifs shared between sequences. We trained SHARK-dive, a homology classifier on a curated set of unalignable homologous sequences enriched in disordered regions. On these sequences, SHARK-dive outperforms conventional alignment methods (SW^6^, BLAST^7^, HMMER^9^) in homology detection and complements recent deep learning based methods (DEDAL). Furthermore, SHARK-dive distinguishes IDR homologs from functionally unrelated sequences reported in the literature. SHARK-dive not only offers predictions of homologous IDRs but also highlights the sequence properties and regions that may contribute to homology. Our tools can be used to facilitate systematic assessment of homology in difficult to align sequences and generate hypotheses of sequence-function relationships of unalignable, disordered sequences.

## Results

### Unalignability and disorder are common in the protein universe

In order to systematically examine the limitations of sequence alignment, we assessed the prevalence of disordered and unalignable regions across Swiss-Prot sequences. Using the disorder predictor IUPred2A^37–39^, we found that 21% of the ∼205 million residues in Swiss-Prot were predicted to be disordered (Fig. 1b) and over 60% of proteins contain an IDR at least 10 amino acids long (Fig. S1), indicating that disorder is pervasive. To quantify the degree to which sequences are alignable, we defined unalignable sequences as difficult-to-align regions which do not map to domains according to Pfam^10^. Despite extensive curation of Pfam over the years, we found that a third of all residues still lack Pfam annotation and thus considered unalignable (Fig. 1b).

To identify relationships between disorder and alignability, we annotated every residue as either ordered/disordered and aligned/unalignable. Whereas only 14% of aligned domain residues are disordered, a far higher fraction (34%) of unalignable residues is disordered (Fig. 1b). Interestingly, we also identified a significant fraction of ordered residues which were not mapped to domains, implying that even some ordered regions are also not amenable to alignment for homology assessment and functional annotation. The poor alignability of IDRs is further highlighted by the low number of disordered seed sequences used to build Pfam families (Fig. 1c). Altogether, this indicates that databases such as Pfam cannot completely categorize and functionally annotate sequences, particularly IDRs, which we reasoned is due to the aforementioned limitations of alignment.

### SHARK-scores are alignment-free, physicochemistry-encoded parameters to compare sequences

In order to overcome the fundamental constraints of alignment, Similarity/Homology Assessment by Relating K-mers (SHARK) was developed to compare sequences in an alignment-free manner (Fig. 2a). Consistent with existing word-based approaches, SHARK first decomposes each sequence into overlapping subsequences of a particular length *k*, which we refer to as *k*-mers (also known as *l*-tuples/*n*-grams/words). Whereas *k*-mer based approaches have been long-established^40–42^, they only recognize identical *k*-mers. Here, we enable the comparison of non-identical but sufficiently similar *k*-mers by assessing their physicochemical properties. According to the Grantham distance matrix^43^, which considers side chain chemical composition, polarity and size, the similarity between *k*-mers is calculated as the mean similarity between amino acids at each position. A *k*-mer similarity matrix is then constructed by comparing all *k*-mer pairs. Aggregation of the matrix yields a SHARK-score representing the similarity between the sequences, of which there are two variants: SHARK-score (best) considers only the most similar *k*-mer, whereas the SHARK-score (*T*) variant considers all similar *k*-mers above a threshold *T* (see Methods). An initial proof-of-concept benchmark revealed that SHARK-scores were superior to local alignment as well as several alignment-free metrics at distinguishing between the functionally opposing N- and C-terminal IDRs of Ded1p orthologs (Fig. S2).

**Figure 2.**
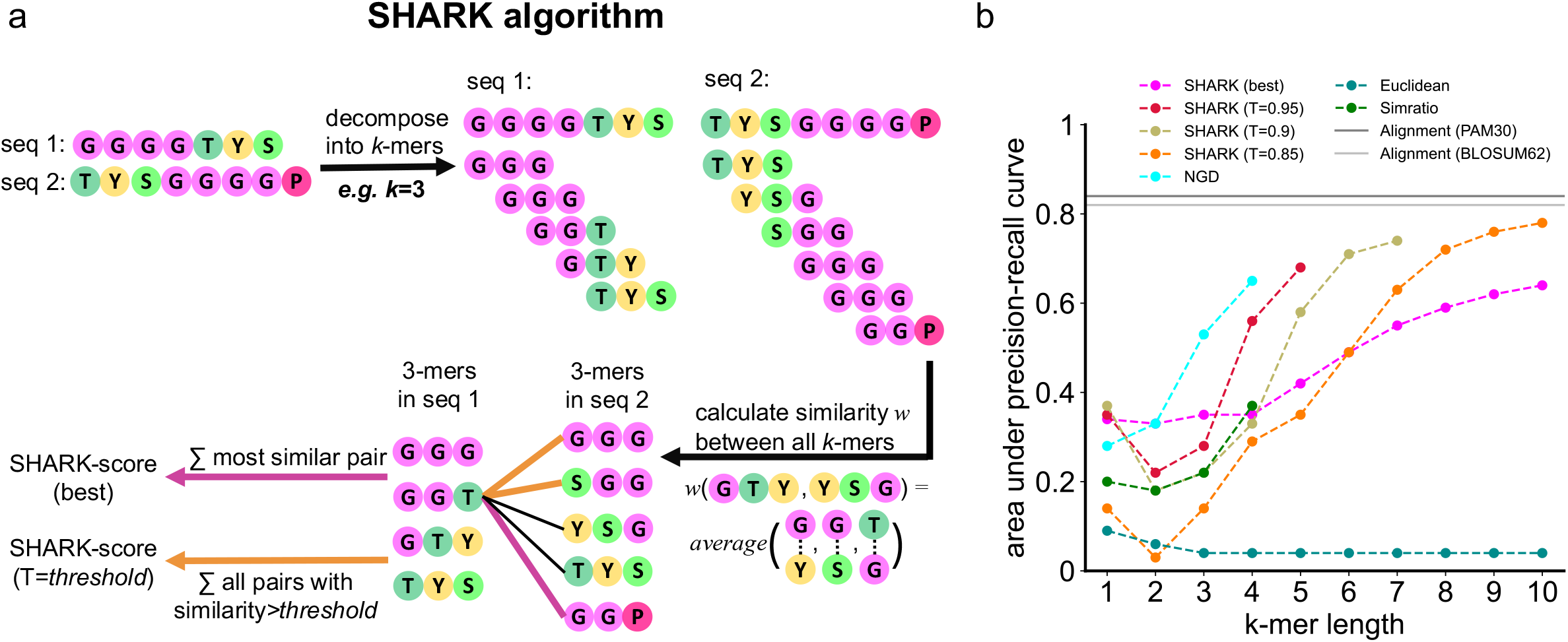
Overview of the SHARK algorithm and SHARK-scores. **a**. SHARK assesses sequence similarity by representing each sequence as subsequences of length *k* (*k*-mers), then assessing the *k-*mer similarities between the sequences. Either the most similar *k-*mer is chosen to give the SHARK-score (best) variant, or all *k*-mers with similarity (*w*) greater than a threshold (T) are summed to give the SHARK-score (T). Unlike most alignment-free scores where a low value indicates higher similarity, a high SHARK-score between sequences indicates high similarity. **b**. Homology detection performance on the alignable-disorder dataset. SHARK outperforms existing alignment-free metrics in homology assessment performance, especially with longer *k*-mers where existing metrics lose performance. SHARK-scores achieve auPRC values greater than 0.65, which is the highest performance attained by existing alignment-free methods (Normalized Google Distance, NGD, at *k*=4). For each algorithm, only *k*-mer lengths where the auPRC would not be over-estimated due to the lack of similar/identical *k*-mers between homologs were plotted (see Methods, Fig. S15). Nonetheless, local alignment still offers superior performance (see Table S1 for details on parameters used).

### SHARK-scores outperform existing alignment-free metrics in alignable IDRs

To assess the efficacy of SHARK-scores in homology assessment, we calculated SHARK-scores on a comprehensive set of homologous IDRs. We extracted a set of 143 Pfam protein families with the highest overall disorder content, henceforth known as the alignable-disorder dataset (Fig. S3, Dataset S1), and performed all-versus-all comparisons between sequences. Due to the class imbalance from the all-versus-all comparison, the overall performance is reported as the integrated area under the precision-recall curve (auPRC). SHARK-scores outperformed existing alignment-free tools tested in terms of highest auPRC attained, afforded by consistently superior performance for longer *k*-mers (Fig. 2b). Whilst SW local alignment was still superior (Fig. 2b), the performance improvements of SHARK-scores suggests that they may complement existing alignment-free algorithms, particularly at longer *k*-mer lengths where existing methods consistently struggled.

### SHARK-dive is a simple machine-learning model trained on a set of unalignable, orthologous sequences

Despite the improvements of SHARK-scores over existing alignment-free metrics in IDR homology assessment, alignment-free approaches were inferior to alignment in the alignable-disorder dataset. We reasoned that since the ground-truth of homology for the dataset is ultimately defined from alignment-based HMMs, this inherently advantaged local alignment in homology assessment.

Accordingly, instead of benchmarking against alignable sequences, we aimed to investigate the performance on homologous but unalignable sequences. However, a large-scale reliable ground-truth dataset was lacking. We therefore adopted an evolutionary-based approach, first using the DisProt database^44^ to identify proteins that contained IDRs with annotated functions which are likely under selection to conserve functional features/motifs, then finding their orthologs using the OMA orthology database (Fig. 3a)^45^. Subsequently, we could assign correspondence between non-domain segments for orthologous proteins with an identical domain architecture solely based on their surrounding domains, without performing sequence alignments (Fig. 3a). A set of these segments was termed a “sequence family”, and all sequence families were aggregated to form the unalignable-ortholog dataset (Dataset S2). We note that although there is no explicit disorder requirement for the orthologous sequences, the majority (70%) of these sequences were indeed disordered (Fig. S4), substantiating the difficulty in aligning IDRs.

**Figure 3.**
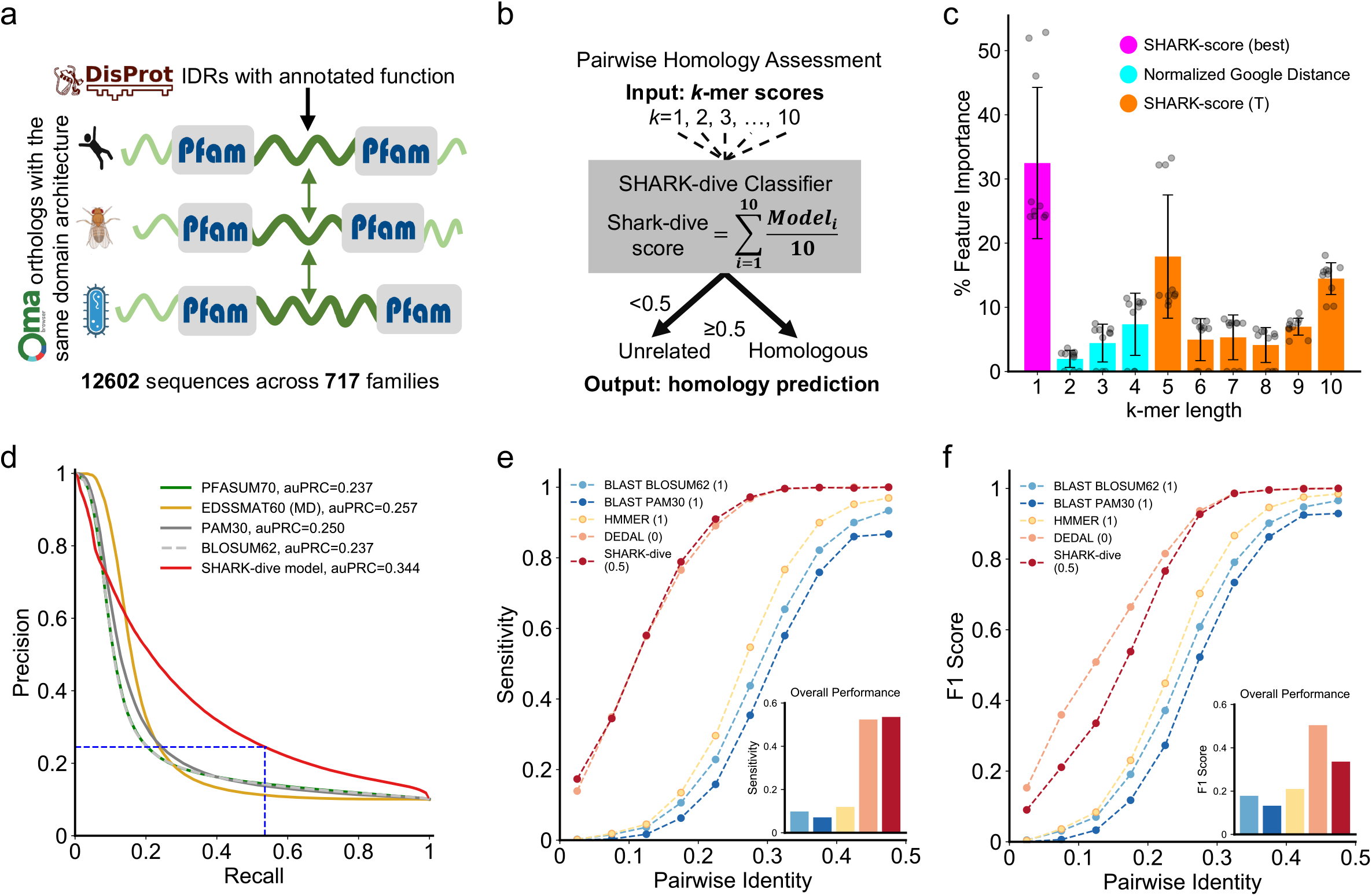
SHARK-dive outperforms conventional alignment in assessing homology in unalignable, disordered sequences. **a**. Curation of the unalignable orthologs dataset. We collected sequence regions that are flanked by the same domains in sequences with identical domain architectures, with functional annotation (from DisProt) for at least one representative ortholog (orthogroups from OMA). Following filtering for max. 50% identity, we curated 12602 unalignable and highly disordered sequences across 717 sequence families. **b**. SHARK-dive is an ensemble gradient boosting classifier of 10 sub-models trained on a training subset of families of the unalignable-ortholog dataset using 10 *k*-mer scores as input. **c**. Feature importance of SHARK-dive. Each bar represents the mean feature importance of the scores for each value of *k* across the sub-models (error bars indicate standard deviation). Magenta and orange bars represent SHARK-scores (best and T variant respectively), cyan bars represent Normalized Google Distance used for *k*=2-4. Grey circles show the feature importance of individual sub-models **d**. Precision-Recall performance of SHARK-dive compared with alignment with various substitution matrices on the unalignable ortholog test sequences. For the EDSSMAT series of matrices only the best-performing one is plotted. Dashed blue lines refer to the performance at 0.5, the default threshold chosen, where SHARK-dive achieves far higher recall at the same level of precision. **e**. Amongst the homology detection tools tested, SHARK-dive offers highest sensitivity to remote homologs across multiple sequence identity (PID) bins. Overall, SHARK-dive offers slight improvements in sensitivity over DEDAL and over four-fold improvement over conventional alignment-based tools BLAST and HMMER (at an E-value threshold of 1, thresholds shown in brackets). Shown in inset is the overall sensitivity across all PID bins. **f**. SHARK-dive and DEDAL offer superior remote homology detection performance over conventional alignment-based particularly when sequence pairs share very low identity. Shown in inset is the overall F1 performance across all PID bins. Thresholds used for each method are shown in brackets.

Since there was a different best-performing alignment-free algorithm for each *k*-mer length in the alignable-disorder dataset, this hinted at the possibility of improved homology detection performance by combining different algorithms across a range of *k*-mer lengths. Based on performance on the alignable-disorder dataset, we chose the best algorithm for each *k*-mer length from *k*=1 to *k*=10 (Table S2), in accordance with the typical motif lengths in IDRs^32,46^. We then split the unalignable-ortholog dataset was then split by sequence family for model training, validation and testing (Fig. S5, Dataset S2). We adopted a machine-learning approach to integrate information across these scores into a homology classifier and named it SHARK-dive. SHARK-dive was trained on the unalignable-ortholog training dataset as a 10-fold ensemble model (Fig. 3b, see Methods). Interestingly, we observed that whilst each *k*-mer length contributed to model decision making, *k*=1,5 and 10 were consistently the most important features (Fig. 3c).

### SHARK-dive has superior sensitivity to alignment in detecting unalignable, disordered homologs

To quantify the performance of SHARK-dive, we assessed its ability to detect homology from an all-vs-all comparison of unalignable-ortholog test sequences that were withheld from model training. SHARK-dive offered superior overall performance and outperformed SW local alignment across all substitution matrices tested, including the PFASUM70 matrix^47^ which had been shown to be the best performing set of parameters for an alignment task based on Pfam-A domains^30^, as well as the EDSSMAT series of disorder-optimized substitution matrices^48^ (Fig. 3d, Table S3).

Whilst auPRC is useful as indicator of overall performance across all thresholds, a specific threshold is used to assess homology in practice. We therefore assessed the threshold-specific performance of SHARK-dive using the default classifier threshold of 0.5 (the discriminatory power of this threshold is shown in Fig. S8). To compare against local alignment, we identified the alignment score threshold defined by the maximum F1 score attained (Table S3). Despite biasing towards alignment, SHARK-dive achieved a 33% increase over the highest F1 score attained by alignment (EDSSMAT60 (MD)), along with a 32% increase over the most sensitive alignment (PFASUM70, Table S3). We further compared SHARK-dive against the most widely-used homology search tools, BLASTp (BLAST) and pHMMER (HMMER). Using default search parameters and reporting thresholds, these tools were not incapable of reporting even 25% of homologs (Table S5). On the other hand, SHARK-dive surpassed the sensitivity bottleneck with a 145% increase the number of true homologs detected (Fig. 3e, inset).

In light of recent advances in applying deep learning and natural language processing models to sequence alignment, we compared SHARK-dive against DEDAL, a novel state-of-the-art pairwise sequence alignment algorithm and homology classifier^30^. DEDAL showed improved performance in homology detection on the test sequences (F1 score=0.505) driven largely by an increase in precision over SHARK-dive (Table S5) whilst maintaining similar levels of sensitivity (Fig. 3e). Nonetheless, SHARK-dive still offers highest sensitivity to remote unaligned homologs across most pairwise sequence identities (PID), whilst maintaining comparable F1 performance to DEDAL (Fig. 3f). Both SHARK-dive and DEDAL offered elevated performance to conventional homology assessment tools such as BLAST and HMMER (Fig. 3e-f) even with more lenient E-value thresholds (Fig. S9), with striking improvements particularly between homologs with lowest sequence identities.

### SHARK-dive identifies unalignable but functionally homologous IDRs

To further validate SHARK-dive, we investigated its ability to identify functionally homologous IDRs reported in the literature. Specifically, we searched for functional rescue experiments reported where protein function was assayed following the replacement of an endogenous IDR (Fig. 4a) with low sequence identity (Table S6), poorly aligned IDRs (Fig. S10). We then assessed if SHARK-dive and existing tools can identify IDRs capable of functional rescue to be homologous. Importantly, we have ascertained that the sequences were not used for model training.

**Figure 4.**
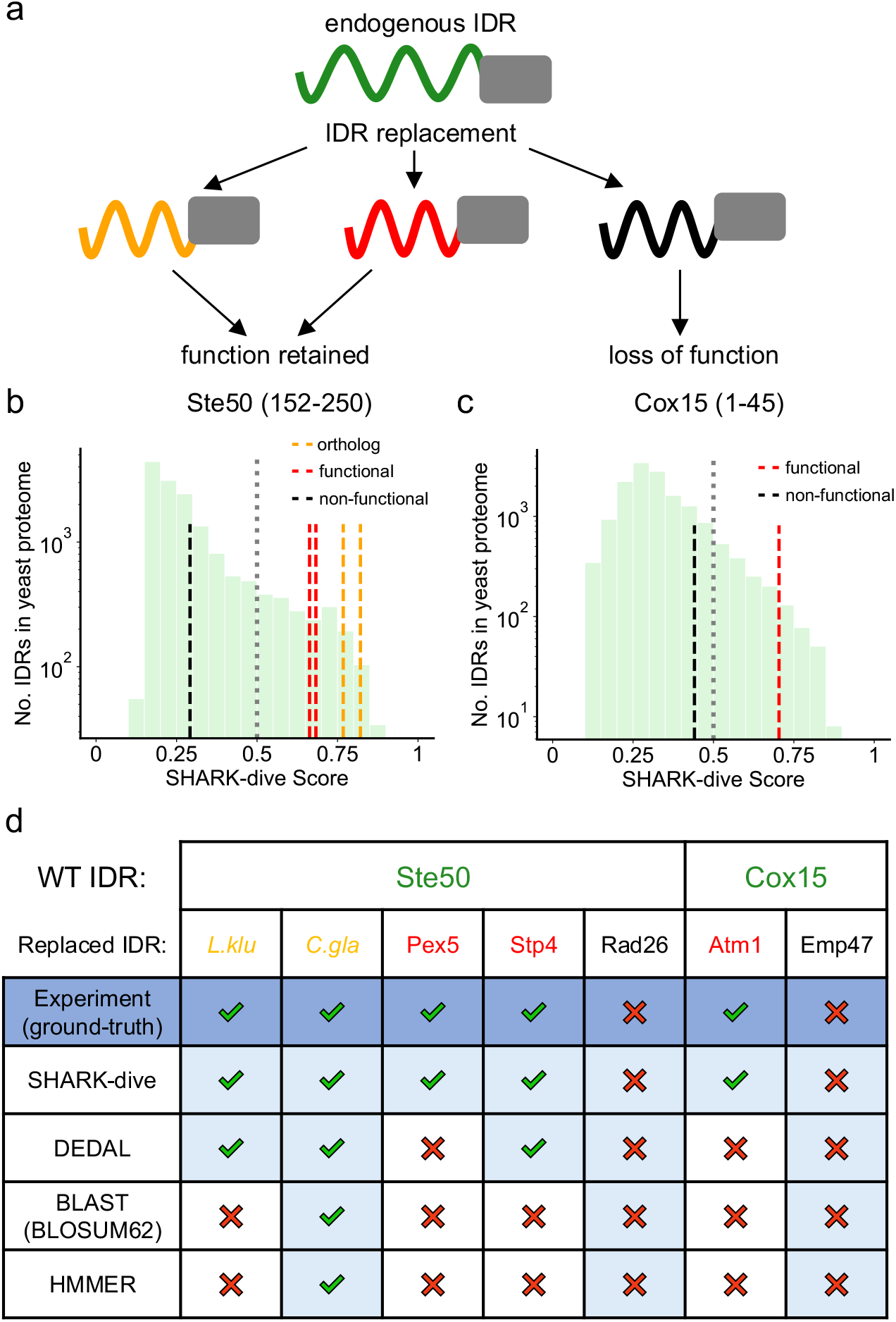
SHARK-dive accurately identifies functional IDR homologs reported in the literature. **a**. Schematic of IDR replacement experiments, where the endogenous IDR (green) is replaced by different IDRs. Some IDRs, including IDRs in orthologs, can rescue protein function (red and orange respectively), whereas others cannot (black). **b**. Highly dissimilar IDRs can rescue Ste50 function upon replacement of its endogenous IDR. IDRs in Ste50 yeast orthologs (orange lines) were able to confer Ste50 WT-fitness and were also predicted as homologous^49^. Pex5 and Stp4 IDRs capable of functional rescue (red lines) were predicted to be homologous to Ste50 IDR, whereas the loss-of-function inducing Rad26 IDR was not^48^. Further, SHARK-dive predicts other IDRs in the yeast proteome (green bars) that may also be homologous to the Ste50 IDR (IDRs above the SHARK-dive threshold, grey line). **c**. Cox15 localization can be rescued upon replacement of its endogenous IDR by Atm1 IDR. SHARK-dive correctly predicts Atm1 IDR to be homologous, whilst the Emp47 IDR was not^48^. Similarly, other IDRs in the proteome are also predicted to be similar. **d**. Summary of prediction accuracy by SHARK-dive and other homology assessment tools on Ste50 and Cox15 IDR homologs, as determined by experimental results. Light blue cells indicate consistent homology predictions to experimental results (dark blue).

Recent work by Zarin *et al*. investigated the effect of replacing the IDR of *S. cerevisiae* Ste50, a mating pathway adaptor^49^. Replacing the endogenous IDR with orthologous IDRs from other yeast species was sufficient to maintain WT-like fitness^49^. SHARK-dive not only correctly identified the homology relationship for orthologous IDRs (Fig. 4b, orange lines), but was able to distinguish *S. cerevisiae* IDRs capable of functional rescue (Pex5 and Stp4, red lines) from the Rad26 IDR which was incapable of functional rescue (black line)^50^. Contrastingly, both HMMER and BLAST failed to detect all functional homologs. Both tools detected only the *Candida glabrata* ortholog below the 0.05 threshold (see Methods), and even with a high E-value threshold of 10 BLAST could not detect homology to the Stp4 IDR. Similarly, DEDAL captured the relationship with orthologous IDRs as well as Stp4, but could not detect the functional homology between Ste50 and Pex5 IDRs (Fig. 4d).

In another set of experiments, mitochondrial localization was restored upon replacing N-terminal IDR of *S. cerevisiae* cytochrome oxidase Cox15 by the Atm1 IDR^50^. This homology was correctly predicted by SHARK-dive, whereas the Emp47 IDR, which could not restore mitochondrial localization, was correctly predicted not to be homologous (Fig. 4c). Neither BLAST, HMMER nor DEDAL could detect Cox15-Atm1 IDR homology (Fig. 4d). Interestingly, both SHARK-dive and DEDAL was capable of identifying synthetic IDRs, designed to share similar molecular/biochemical properties as Cox15 IDR, and also capable of conferring mitochondrial localization upon replacement^51^ (Table S8). Altogether, SHARK-dive accurately predicted functional homology of IDRs across different functions/phenotypes despite negligible sequence similarity.

### SHARK-dive offers interpretable predictions of homologs

Given SHARK-dive’s accuracy in identifying experimentally-verified functional IDR homologs, we used SHARK-dive to generate predictions of homologous IDRs. We used the human FUS as an example, that has been shown to form *in vitro* condensates^35^ as well as been recruited to stress granules *in vivo*^52,53^. Specifically, the disordered N-terminal prion-like disordered region of human FUS (FUS PLD, 1-211, Fig. 5a) mediates condensate formation via homotypic Tyr-Tyr π-π interactions^54^, or Tyr-Arg π-π and Arg-Tyr cation-π interactions with its highly disordered C-terminal arginine-rich RNA binding domain (FUS RBD, defined as residues 212-526, although it contains two Pfam domains, Fig. 5a)^35^.

**Figure 5.**
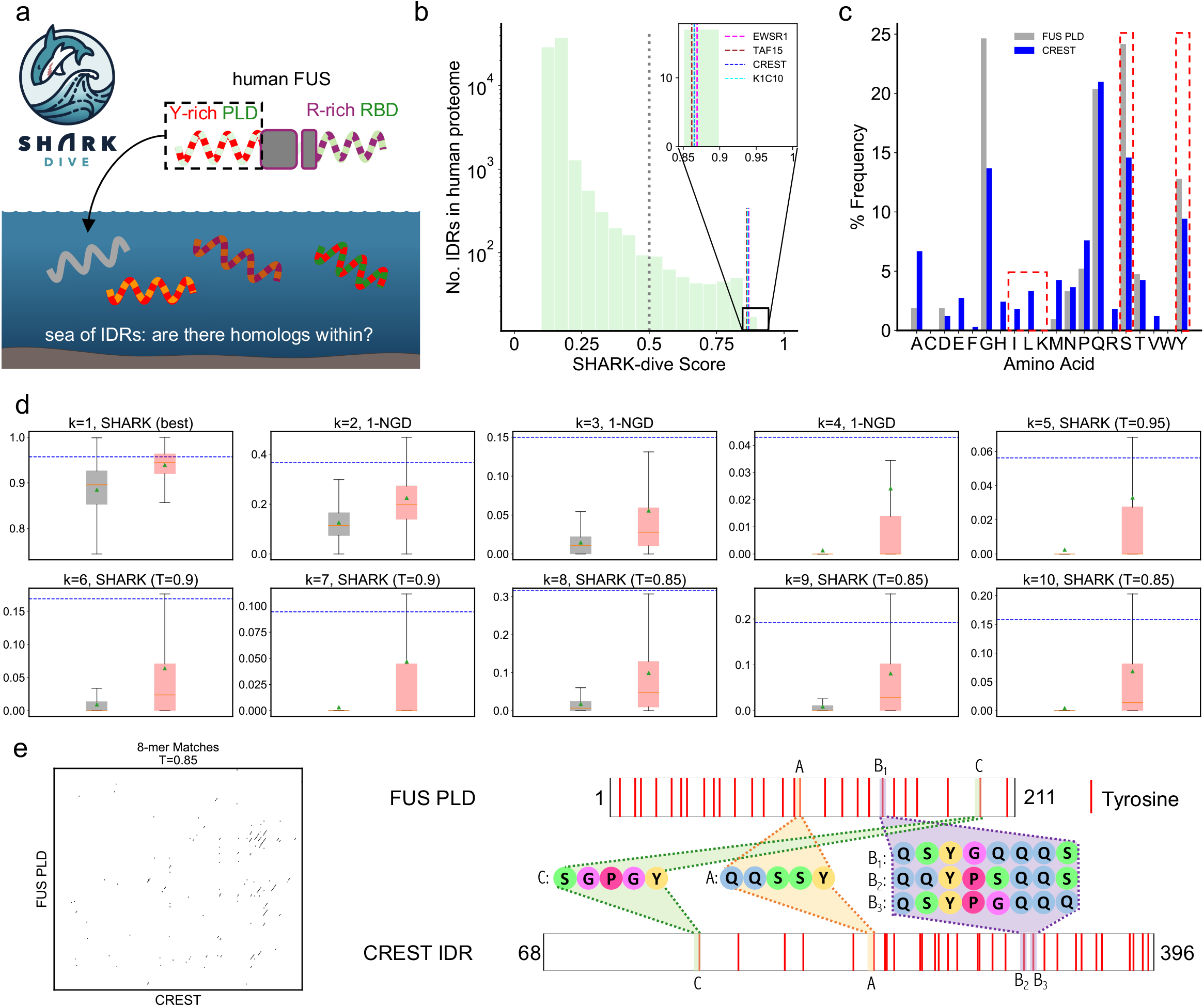
SHARK-dive offers interpretable predictions of homologous IDRs. **a**. Schematic of the homology search with human FUS PLD (aa 1-211 as defined in Wang *et al*. 2018^35^) to find similar IDRs. **b**. SHARK-dive predicts various IDRs in the human proteome to be homologous, including N-terminal IDRs of known FET family members TAF15 and EWSR1, as well as a variety of unrelated IDRs in proteins such as transcriptional activator CREST and Type 1 Keratin (K1C10). **c**. Human CREST IDR and FUS PLD share similar amino acid compositions, particularly in its high tyrosine and serine contents and low abundance of lysine and aliphatic amino acids (highlighted in red). **d**. To interpret the homology prediction of SHARK-dive, the input features (*k*-mer scores) can be compared to the distribution of unrelated sequences or true homologs in the training sequences to identify particular *k*-mer lengths which may contribute to prediction of homology (or lack thereof). In the case of the CREST IDR which is confidently predicted to be homologous, all *k*-mer scores are higher than the average homologous pair. For *k*=2-4, NGD scores are visualized as 1-NGD (otherwise known as the Normalized Google Similarity) such that higher scores represent higher similarity for ease of interpretation. Boxplots show median (orange line), mean (green triangle) and quartiles. Outliers are not shown but can be found in Fig. S11. **e**. The inputs to SHARK-dive are similarity-matrix derived *k*-mer scores, which can be visualized to show regions of high similarity between sequences. This can be visualized for a range of *k***-**mer lengths as well as similarity thresholds (*k*=8 and T=0.85 shown here). Following this, similar *k*-mers identified from the similarity matrix can be mapped back onto the sequence to identify potential motifs or sequence characteristics that may contribute to shared function/homology (3 examples here shown). Potential motifs can be in different positions along the sequence such as motifs A and C. The similarity matrix also allows similar but non-identical motifs can also be identified (e.g. B_1,2,3_) as well as detection of multiple motifs matches (e.g. B_2_ and B_3_ are both similar to B_1_). Inspection of the shared motifs suggests that, besides the tyrosine and serine residues, glutamine blocks (e.g. QQ) may also contribute to shared homology between CREST IDR and FUS PLD.

We hypothesize that sequences homologous to the FUS PLD may be able to interact with FUS to drive phase separation and biomolecular condensate formation, and used SHARK-dive to find its homologs in a set of IDRs extracted from the proteomes of human and other common model organisms (see Methods). SHARK-dive not only predicted homology to the N-terminal IDR of the mouse FUS ortholog, but also to the PLD-containing N-terminal IDRs of FET-family members^55^ TAF-15 and EWSR1 that are known to phase separate^35^ (Fig. 5b). Moreover, SHARK-dive identified several IDRs in other organisms to be similar to the FUS PLD (Fig. S13, Dataset S3).

As an example, SHARK-dive predicted an IDR in CREST to be homologous (Fig. 5b). CREST is a highly disordered transcription activator and regulator of cortical neuron dendritic growth^56,57^. Similar to FUS, it is also implicated in amyotrophic lateral sclerosis^58,59^. It contains a C-terminal IDR that shares only 15% identity with FUS PLD, and not reported in a BLAST search of the FUS PLD (from NCBI BLAST webserver). Importantly, homology predictions of SHARK-dive can be interpreted by looking at signals from its input features, i.e. the *k*-mer scores and the similarity matrices used to derive them. The amino acid composition (*k*=1) shows high similarity between FUS PLD and CREST (Fig. 5c), corroborated by the high 1-mer score which exceeds the mean for true homologs in unalignable-ortholog training sequences (Fig. 5d). Similarly, the 5-mer SHARK score (T=0.95) showed the same behavior, and comparison of CREST *k*-mers revealed correspondence to multiple regions in the FUS PLD (Fig. S12). Interestingly, some of these 5-mers are swapped in order between the two sequences and thus would be difficult for alignment to fully detect (Fig. 5e). As a comparison, a predicted non-homolog ZN365 shows markedly fewer *k*-mer matches and lower scores similar to those of unrelated sequences used to train SHARK-dive (Fig. S14b).

While both CREST and FUS have regulatory roles in transcription and neuronal growth, we also report that the FUS PLD was also predicted to be homologous the C-terminal IDR of human Type I Keratin (K1C10/Krt10), which shares no functional similarity to FUS at all; it is involved in skin epidermal barrier formation^60^. Again, corresponding *k*-mers can be found between FUS and the keratin C-terminal IDR (Fig. S14a). We further note that both FUS^61^ and ZN365^62^ have also been shown to be involved in double-stranded DNA break repair, suggesting that functional similarity of the protein overall is independent of the IDR homology prediction and that SHARK-dive can be used to predict IDR homology independent of the function of the full-length protein.

Compellingly, literature review revealed that these IDRs have indirect experimental support for their homology to the FUS PLD. Like FUS PLD, CREST has already been reported to be capable of phase-separation and in fact can even interact with the FUS PLD^58,59^. We note that homology was not predicted between FUS RBR and the CREST IDR, suggesting that the prediction is specific to the IDR and not due to the protein overall. Krt10 has also been shown to phase separate promote liquid-like keratohyalin granules via its N- and C-terminal low complexity domains^60^. Coupled with the ability to determine the regions between sequences that contribute to their predicted similarity, we believe SHARK-dive offers the ability to generate hypotheses regarding IDR homology and function across proteomes and species.

## Discussion

Alignment-based homology detection has exponentiated our understanding of sequence-function relationships and aided functional annotation in structured regions but has had limited benefit for disordered and rapidly evolving regions. Here, we present the SHARK algorithm and its derivatives SHARK-scores and SHARK-dive, developed specifically to tackle this challenge. It is based on an alignment-free, *k*-mer based approach which breaks free from the inherent constraints and limitations of sequence alignment, and innovates by assessing the similarity between *k*-mers in physicochemical space. The SHARK algorithm amalgamates the strengths of both alignment and alignment-free approaches by combining the removal of collinearity constraints with the consideration of *k*-mer similarity.

Despite the improvement of individual SHARK-scores over existing alignment-free algorithms, we reasoned that accurate homology detection for disordered, unalignable regions requires consideration of amino acid composition as well as *k*-mers of varying lengths. Besides the performance gain in detecting remote homologs, this was also biologically motivated since functional motifs may be of varying sizes. We therefore harnessed the power of machine-learning in aggregating a range of *k-*mer scores to train the SHARK-dive model. We note that SHARK-dive is a classification model, thus its output is not the similarity between the two sequences but rather the probability that they are homologous. In the future, it would be interesting to extend SHARK to compare multiple sequences and have an iterative search approach of large databases analogous to alignment-based search algorithms such as PSI-BLAST^7^ and JackHMMER^63^ to further increase sensitivity to remote homologs.

Since previous remote homology assessment benchmarking datasets were based on ordered/structured regions (e.g. SCOPe, ASTRAL)^42^ or alignable sequences (e.g. Pfam)^30^, here we curated a dataset of unalignable orthologous sequences enriched in IDRs. We identified non-domain sequences between known orthologs that share identical domain architectures (Fig. 3a) and are flanked by the same set of domains. Our domain-based approach is an alignment-independent extension of the protocol used by Lu *et al*. and similar to approaches adopted in experimental contexts^49,64,65^. There are nonetheless limitations of our approach, such as the assumption of non-homology between different sequence families, which is difficult to guarantee without extensive experimental evidence. This is nonetheless a general assumption held also by existing benchmarks, which we aimed to alleviate with the strict identity filtering (<50% identity) applied to our sequences (see Methods).

Curation of the unalignable-ortholog dataset also enabled a systematic analysis of the suitability of existing alignment tools in detecting relationships in these difficult-to-align sequences. We report that SHARK-dive outperforms widely used alignment algorithms such as Smith-Waterman, BLAST and HMMER in finding homologs, and performs on par with a recent deep learning and language model-based tool, DEDAL. DEDAL was so far only evaluated on a homology detection task of Pfam domain sequences, here we show that it also performs excellently on non-domain regions with better accuracy and slightly lower specificity than SHARK-dive (Figs. 3e,f).

We wish to highlight the complementary strengths of SHARK-dive and DEDAL. We opine that the excellent homology detection performance of highly advanced, complex deep learning methods such as DEDAL comes at the cost of the interpretability of its predictions. In contrast, SHARK-dive is a relatively simple, gradient-boosted decision tree classifier based on only 10 sequence features (i.e. *k*-mer scores). Therefore, the rationale behind SHARK-dive predictions is more interpretable. To demonstrate this, SHARK-dive was used to predict homology between the highly disordered FUS PLD and sets of IDRs in different proteomes (Fig. 5a). Several examples of seemingly unrelated IDRs were investigated and their similarities to the FUS PLD annotated and visualized (Figs. 5e, S12-14). Coupled with the (albeit indirect) support of literature evidence of their similarity, we surmise that SHARK-dive offers tractable hypotheses of the sequence regions and properties that may underlie their functional homology. This can then be further investigated in the future by IDR replacement experiments with predicted homologs (Fig. 4a)^49,50^, followed by mutations of these predicted motifs or compositional characteristics to ascertain their necessity and sufficiency in driving protein function. We also note that whilst these analyses focused on IDRs, SHARK-dive should be detect homology not only between IDRs but all difficult-to-align sequences.

To that end, the superior sensitivity of SHARK-dive, combined with its less demanding computational requirements, make it a convenient option for quick searches of IDR homology. For more extensive studies, we propose that a pipeline consisting of homology prediction by a general purpose tool such as DEDAL to identify regions of significant and extensive sequence similarity between IDRs that could yield a high-quality alignment. If alignment is poor, the more specifically-tuned, alignment-free

SHARK-dive could be applied systematically and proteome-wide to facilitate the study of sequence-function relationships in disordered, difficult-to-align regions.

## Acknowledgements

This project was funded by the Max Planck Gesellschaft. CFWC was supported by the Deutsche Forschungsgemeinschaft (DFG, German Research Foundation) under Germany’s Excellence Strategy – EXC-2068 – 390729961– Cluster of Excellence Physics of Life of TU Dresden. AH was supported by the ELBE postdoctoral fellowship. We thank Cedric Landerer, Simon Alberti and Tony Hyman for their valuable insights during the development of the project. We would like to thank the Computer Services and Scientific Computing Facilities of the MPI-CBG for their support, especially to Oscar Gonzales for supporting our HPC. We thank Michele Marass for the feedback on the manuscript and Hannah Jor for the design of the logo.

## Data and code availability

The code base and readme files can be found at https://git.mpi-cbg.de/tothpetroczylab/shark. Further, we provide jupyter notebooks to aid easy interpretation and visualization of SHARK-dive scores: https://git.mpi-cbg.de/tothpetroczylab/shark/-/tree/master/notebooks.

## Author Contributions Statement

ATP and CFWC conceived the project. CFWC performed research, and algorithm development. SG designed and implemented the software. CFWC prepared figures. CFWC, AAH and ATP analyzed data. ATP and CFWC wrote the manuscript with inputs from all authors.

## Online Methods

### UniProt-wide analysis of disorder and definition of alignability

All 568002 Swiss-Prot sequences in the 2022-03 release of UniProt were retrieved^2^. For each sequence, we mapped the domains (defined by the alignment coordinates to the profile HMM) using HMMscan using the Pfam35.0 database with the gathering thresholds as a cutoff^10^. Sequences not longer than 10 residues were removed. Each residue was then annotated as ordered/disordered or domain/non-domain depending on its IUPred2A disorder score (long option)^37,39^, and its Pfam annotation respectively. We use a threshold of 0.4 to account for disordered regions that have a propensity for disorder-to-order transitions, and define unalignable regions as those that do not map to Pfam domains (non-domain).

### Extraction of IDRs

Reference proteomes were obtained from the 2022-05 release of UniProt for 7 organisms: *H. sapiens, M. musculus, E. coli, S. cerevisiae, A. thaliana, D. rerio, D. melanogaster* (Table S7). Each sequence was analyzed using the disorder predictor IUPRED2a (long)^37^, with smoothing applied to residue-specific disorder values using a Savitzky-Golay filter (moving average window size=9, polynomial degree=3). Setting the disorder threshold at 0.4, consecutive disordered residues were concatenated into IDR segments. To identify the number of proteins with significant disorder in Swiss-Prot, the length of the longest IDR segment present was chosen. For IDR homology searches in proteomes, only ≥10 amino acid IDR segments without unresolved or non-canonical amino acids were considered.

### Development of the alignment-free SHARK algorithm and SHARK-scores

In summary, SHARK-scores compares sequences using 6 main steps. The first 4 steps form the core of the SHARK algorithm:

1) Each sequence (query *Q* and target *T*) is decomposed into a *k*-mer vector, where *k* represents the length of the peptide. Each *k*-mer vector encodes the identity of the *k*-mer and the frequency (*q*_*i*_ or *t*_*j*_) at which it occurs in the sequence, where *i* and *j* represent each unique *k*-mer in sequence *q* and *t* respectively.

2) The similarity between 2 amino acids (*D*′) is derived from an amino acid physicochemical distance matrix (*G*) (in our case the Grantham’s Distance matrix^43^). To convert from a distance (i.e. a measure of dissimilarity) to a similarity measure, the distance values *D*′ are inverted and min-max normalized to give *k*-mer similarity score *D*:

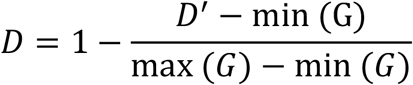

3) Between two *k*-mers *i* and *j* of identical length, the similarity *w* is calculated as

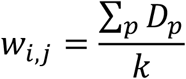

i.e. the mean physicochemical similarity across each amino acid position (*p*) of the *k*-mers. As such, *w*=1 for identical amino acids and *w*=0 for the most dissimilar pair (Cysteine-Tryptophan).

4) Between two sequences, all k-mers are compared against each other to form a similarity matrix (*M*), which gives the *k*-mer similarity (*D*) between the unique *k*-mer pairs of both sequences.

5) For each *k*-mer pair (*i,j*) in *M*:

a) All but the most similar *k*-mer in the other sequence are filtered out (*w* set to -1) to give a sparse matrix. Each non-zero element in the matrix is multiplied by the frequency product in both sequences *q*_*i*_ · *t*_*j*_ (*where w* > −1) and weighted (divided) by a length correction (LD) function, defined as:

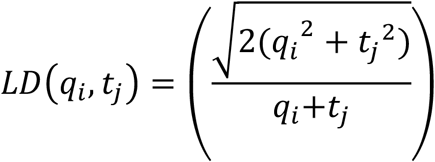

The LD function serves to account for the difference in frequencies between *k*-mers. In the case where multiple *k*-mers have equivalent similarity, the most similar *k*-mer is chosen based on the lowest LD value. Finally, this is summed across the entire matrix and normalized by the sum of the frequency products of non-filtered *k*-mers in both sequences, ∑ *q*_*i*_ · *t*_*j*_ (*where w* > −1), to give the best-match SHARK-score (SHARK-score (best)). Identical sequences will score 1.

b) All *k*-mers not exceeding a similarity threshold (*t*) are filtered out (*w* set to -1) to give a filtered matrix. Resultingly, each *k*-mer (*i* or *j*) contains only sufficiently similar matches to itself, represented by values > -1 in its row/column (row for query *k*-mers *i*, column for target *k*-mers *j*). For each *k*-mer, its row/column score is then aggregated by calculating the weighted average similarity across the row/column, again using LD to factor in the difference in *k*-mer frequencies between the *k*-mer and the total number of matches sufficiently similar *k*-mer matches in the other sequence. In other words, each *k*-mer is compared to a “composite” *k*-mer that represents the overall similarity of the matches:

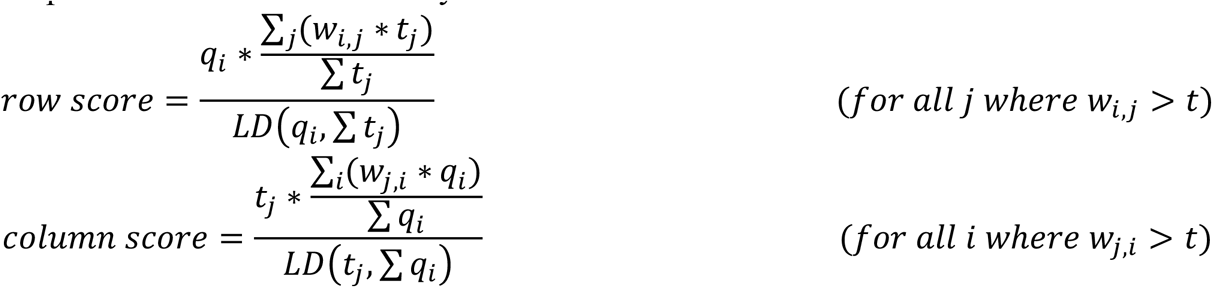

All row and column scores are summed (i.e. for all unique *k*-mers across both sequences) and then normalized by the total number of *k*-mers across both sequences, ∑ *q*_*i*_ + ∑ *t*_*j*_, to give the filtered SHARK-score (**SHARK-score (T=*t***). We note that the filtered score does not guarantee that identical sequences get the maximum score of 1, but is designed to fully capture the correspondences between all sufficiently similar *k*-mers between 2 sequences.

6) The matrix is invariant to which sequence is used as query and target (the similarity matrix *M* is simply transposed), ensuring that scores are symmetrical.

### Curation of disordered Pfam families (alignable-disorder dataset)

Pfam is a repository of homologous domain sequences grouped into families^10^. Sequence homology is assessed by concordance to an alignment-based Hidden Markov Model built from manually-curated seed sequences.

To select for the most disordered Pfam families in Pfam 34.0 (released March 2021)^10^, the disorder in the 90% non-redundant seed sequences of each family was analyzed using IUPRED2a (long option)^37,38^. For each sequence, the mean disorder score across all positions (mean sequence disorder) was calculated; redundant sequences (>90% identity) and sequences where the seed and the full alignment differed were discarded from analysis. For each family of seed sequences, the mean sequence disorder across all seed sequences were averaged to give the mean seed disorder.

1440 Pfam families with mean seed disorder >0.5 were selected for further analysis, and their domain sequences (full alignment) were obtained from Pfam. CD-HIT was applied to filter out sequences with more than 50% identity; families with fewer than two sequences were also ignored since sample standard deviation cannot be calculated. 1374 disordered families remained from the analysis. From these families, the mean sequence disorder was calculated for all domain sequences of that family and averaged to give the mean family disorder; the variation in mean sequence disorder was also calculated as the standard deviation of family disorder. Within these 1374 families, a dataset was curated containing families in the highest 25% of mean family disorder and lowest 25% standard deviation of family disorder, ensuring that most (if not all) sequences are disordered. Finally, sequences shorter than 10 amino acids or containing unresolved amino acids (“X”) were removed, giving 2583 sequences across 143 Pfam families (Dataset S1).

### Curation of unalignable, orthologous sequences from DisProt (unalignable-ortholog dataset)

Entries with functional annotations were curated from the DisProt database (2022-03 release)^44^, this includes all entries with annotated molecular function/disorder functions in the ‘term_namspace’ column. Orthologous proteins across the tree of life were identified using the OMA orthology database (Dec 2021 release)^45^. Orthologs in multiple orthology groups, as well as sequences with ambiguous OMA identifier – UniProtID or sequence – OMA identifier mappings, were excluded.

Using HMMscan against the Pfam35.0 database with the gathering thresholds as a cutoff, we mapped the alignable domains (defined by the alignment coordinates to the profile HMM) onto each sequence to obtain its domain architecture. Domains were mapped sequentially to avoid overlap, with the most significant domains (in ascending order of sequence E-value) mapped first. Subsequent domains that overlap with mapped domains were excluded, ensuring that each residue is only mapped to one domain. Each protein is thus represented by its domain architecture consisting of an N-terminus, followed by all mapped Pfam domains linked by non-domain sequences, and ending with a C-terminus (Fig. 3a). Using the protein entry curated in DisProt as a reference, we selected only for orthologs with an identical domain architecture. This allowed correspondence to be established between each non-domain segment across orthologs, solely based on their surrounding domains. We then mapped the amino acid coordinates (start and end positions) of each functional IDR region onto the sequence. Where the IDR overlaps with a non-domain segment, we extracted the corresponding non-domain segments in orthologs flanked by the same set of domains to give a family of orthologous unaligned segments (henceforth referred to as a sequence family). In rare cases (0.26% of sequences in the final dataset across 2.2% of sequence families) where the DisProt-annotated IDR regions of the reference entry are fully mapped by Pfam domains, non-Pfam annotated regions in orthologous sequences were still extracted since they still correspond to the same sequence family (flanked by the same set of domains). In cases where the DisProt function annotation spans multiple non-domain regions, we extracted each sequence family separately.

All sequence families were aggregated, segments not longer than 10 amino acids or containing unresolved amino acids were removed, and filtered for <50% identity with CD-HIT^66^. Due to the requirement during our dataset curation that Pfam domain regions must be non-overlapping and sequentially mapped by sequence E-value, we performed a final HMMscan on sequences and removed 183 sequences (1.4%) that contained Pfam domains, ensuring that our sequences are fully non-Pfam aligned. This yielded a set of 12602 sequences across 717 families of orthologous unaligned segments (Dataset S2).

### Ded1p orthologous N- and C-termini sequences (Ded1p dataset)

Eukaryotic orthologs of *S. cerevisiae* Ded1p (UniProt ID P06634) were extracted using the EGGNOG orthology database (v5.0). From the multiple sequence alignment of the orthologs with MUSCLE (v3.8.31), the N- and C-termini IDRs were obtained according the helicase core domain boundaries of the *S. cerevisiae* sequence, defined as positions 98-535 according to alignment to the human ortholog DDX3X for which a crystal structure is available (PDB 2I4I, 4PXA, 5E7I). The MSA was manually curated and removal of redundant, highly similar (>95% identity) sequences was performed using CD-HIT. This gave a set of 268 orthologs each with an N- and C-terminal IDR (536 sequences in total).

### Performance of homology assessment

All-vs-all sequence comparisons were performed and the ability of the various algorithms to classify homologs was assessed by plotting the receiver operating characteristic (ROC) and precision-recall (PRC) curves.

In the case where the number of true positive (homologous, TP) and true negative (unrelated, TN) sequence pairs are similar such as the Ded1p dataset, the auROC provides a more balanced measure. For the alignable-disorder and unalignable-ortholog test datasets the overall performance is reported as the integrated area under the precision-recall curve (auPRC). This is due to the class imbalance from the all-versus-all comparison, which yields a far greater number of unrelated sequence pairs than true homologs. The auPRC quantifies the ability to detect true positives whilst minimizing false positives in the prediction and is more suitable in situations with a class-imbalance.

Besides SHARK scores and the SHARK-dive model, the following alignment-free metrics were also compared: Normalised Google Distance (NGD)^67^, Similarity Ratio (Simratio)^68^ and Euclidean distance. NGD and Simratio was reported as the best performing metrics in an alignment-free homology assessment benchmark^42^, whilst Euclidean Distance is a commonly used distance metric. The alignment-based algorithms HMMER, BLAST and DEDAL and Smith-Waterman local alignments (with Biopython pairwise2 module) were also benchmarked.

Alignment-free scores can reach maximum dissimilarity and lose sensitivity to remote homologs with longer *k*-mers because there are no similar/identical *k*-mers shared between homologs. Accordingly, we define the ‘area of non-sensitivity’ as the region in which the scores are unable to distinguish between true homologs and unrelated sequences, calculated via the trapezium rule between the final recall point (recall=1.0) and the previous recall point. For the alignable-disorder dataset, a threshold of 0.003 was set such that only *k*-mer lengths and scores with area of sensitivity< 0.003 are considered suitable and subsequently used, so as not to over-estimate their precision-recall performance (Fig. S15).

Protein-protein BLAST (BLASTp) was performed using both standard (blastp) and short (blastp-short) options. Sequences databases were generated from FASTA files using the makeblastdb command, which was then used to search against by query sequences (in FASTA files). To maximize sensitivity and detection of remote homologs the E-value threshold was set to 10^4^. Default parameters were used for BLAST searches, and we also included the following substitution matrices and gap penalties (Table S). Note, that we use BLAST’s definition of gap cost. Specifically, given a gap of length *n*:

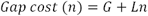

where G is the gap opening cost and L is the gap extension cost.

For HMMER the reporting and inclusion E-value thresholds (-E and -incE) were set to 10^14^ once again to maximize sensitivity, alongside the --max option that turns off sequence filters and runs the full postprocessing on all targets to ensure maximum sensitivity.

Unless otherwise specified, local alignment was performed using Smith-Waterman dynamic programming with the indicated substitution matrix with parameters in accordance with default values and definition of gap costs used in BLASTp searches (Table S1).

DEDAL homology detection/classifier tool was used, which outputs homology detection logits (log-odds score). A threshold of 0 is used, corresponding to a probability of homology =0.5.

### SHARK-dive model feature selection and training

Feature selection of the SHARK-dive model was based on the performance of individual scores on the alignable disorder dataset. We selected 10 features, one scoring method per *k*-mer length from 1-10 (Table S2).

A 70:20:10 train-validation-test split of the unalignable ortholog dataset by family was performed. We split sequences at the family level to ensure that test dataset sequences are not homologous to sequences used in training. We further split the validation set in half for early stopping to prevent overfitting (validation 1) and hyperparameter tuning (validation 2), respectively. We verified that IDRs were retained in all splits (Fig. S5) during the train-test split. The final numbers of sequences and sequence families in each split are detailed in Fig. S5e.

To account for the variability in family size and sequence lengths, we developed a 10-fold ensemble model where each model is trained on a particular subset of families. Accordingly, the training dataset was split into 10 overlapping folds by orthologous segment family. Each fold was held out once for cross-validation (cross-validation fold), and the 9 remaining folds combined into a training fold used to train the model. Accordingly, this gives 10 sub-models, each trained on a large training fold from the combination of the 9 folds with the stated hyperparameters. The validation1 sequences were introduced as an early-stopping dataset to prevent overfitting, with training stopped when the performance on the validation1 sequences does not increase in the following 100 rounds. All training and cross-validation folds, as well as the early-stopping validation1 dataset, were balanced for number of homologs and number of unrelated sequences (same number in of TP and TN). We verified that the choice of TN does not affect model performance (Fig. S6a,b). To tune the CatBoost^69^ model hyperparameters, we tested 250 combinations of three hyperparameters via a grid search (Dataset S4) and then selected the best performing model based on F1-score on the validation2 dataset. This corresponded to a number of weak learners of 700, maximum depth of 8 and learning rate of 0.1. The cross-validation performance of each sub-model (i.e. each training fold) on its corresponding validation fold is shown in Figure S7. We further note that the feature importance of each sub-model has similar trends (Fig. S6c). The final SHARK-dive score is the unweighted mean prediction across all 10 sub-models.

### Percentage Sequence Identity (PID) calculation

Sequences were globally aligned using Biopython pairwise2 module, with BLOSUM62 matrix and standard gap penalties (Table S1). End gaps were not penalized. Percentage sequence identity was defined as fraction of identical residues in the alignment (aligned residues + gaps within the alignment i.e. the length of the alignment), i.e.

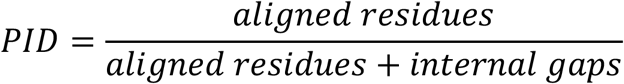

and lies in accordance with the Doolittle definition^70,71^.

### Performance evaluation

For local alignment scores with Smith-Waterman, the threshold of homology was chosen based on the optimal F1 score on the withheld test set, specific to each substitution matrix and gap penalty parameters. Threshold for existing alignment-based homology search tools (BLASTp, pHMMER) was chosen based on default E-values and bit-scores used. Performance of stricter (i.e. lower) E-values are also reported (Table S4). Following selection of the threshold, threshold-specific performance metrics such as sensitivity/recall, specificity, accuracy and F1 were also calculated (Table S5). PID-stratified sensitivity and F1 performance is also reported.

### IDR replacement/functional rescue validation

Sequences were obtained from the publications reporting such IDR replacement experiments, directly from supplementary information or from Yeast Genome Browser^72^-UniProtID mappings^49,50^. For all tools SHARK-dive, DEDAL, pHMMER and BLAST (BLOSUM62), a database comprising all replacement IDRs as well as the WT IDR was curated, and a WT-versus-all comparison performed. To account for the smaller size of the sequence database against which sequences are searched (<10 sequences versus the ∼1800 sequences for the unalignable-ortholog test dataset), E-values for homology detection are accordingly adjusted to 0.05 since they depend on the size of the database (we adopt BLAST’s definition of database size as the number of residues in the database^73^). Multiple sequence alignments were performed using MUSCLE (v3.8.31) unless otherwise indicated^74,75^.

