## Supplementary Information for "SHARK enables homology assessment in unalignable and disordered sequences"

Affiliations

Supplementary Figures 1-15

Supplementary Tables 1-9

Supplementary Datasets 1-4

Supplementary References

### Supplementary Figures

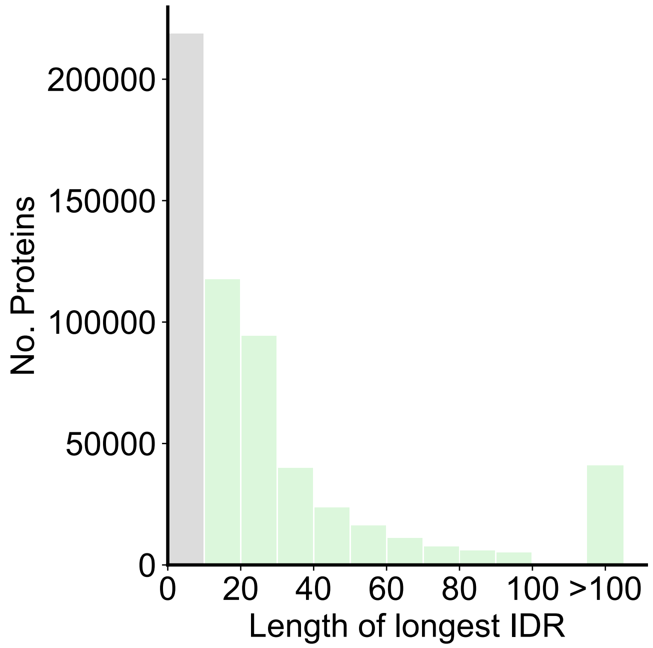


**Figure S1. Most proteins contain a disordered region.** Distribution of the longest IDR found in each curated Swiss-Prot protein (*n*=567,192). 63.7% of proteins contain an IDR of at least 10 amino acids in length (green).

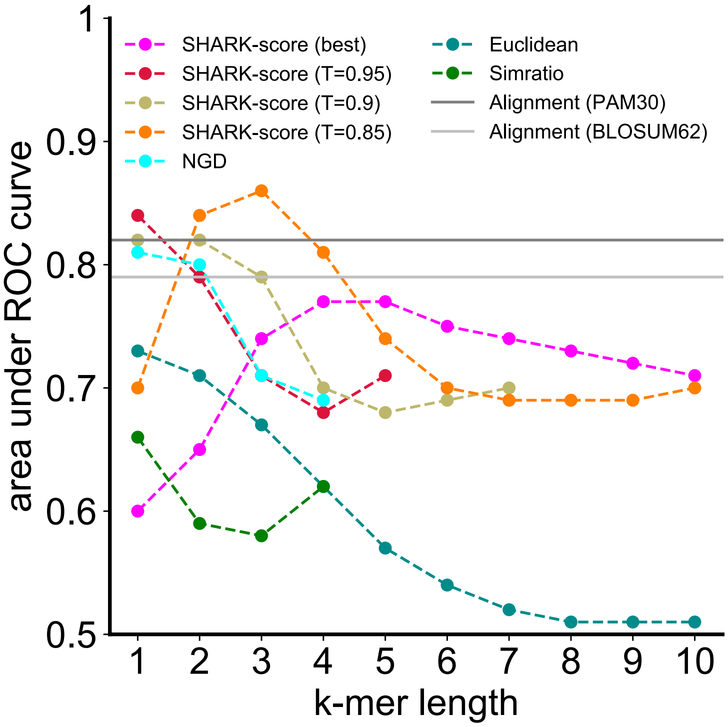


**Figure S2. For a set of Ded1p orthologs, SHARK-scores outperform alignment and existing alignment-free metrics in differentiating sequences which have opposing functional effects.** In order to assess the ability of SHARK-scores in identifying functionally similar sequences, we performed a proof-of-concept test on a set of eukaryotic Ded1p orthologs. Work by Iserman *et al.* identified opposing functions of the N- and C- terminal IDRs on phase separation behavior in Ded1 (Fig. 1a)[1]. Specifically, the N-terminal IDR inhibits heat-induced phase separation of Ded1 whilst the C-terminal promotes condensate formation via protein-protein interactions. Since these functions are likely under selection and thus conserved between orthologs, a set of Ded1p N- and C- terminal IDR orthologs were curated.

We assessed the ability of different algorithms to capture the homology between IDRs from the same terminus and observed that alignment-free metrics generally outperform Smith-Waterman dynamic-programming algorithm (henceforth referred to as local alignment). In addition to the commonly used Euclidean distance, we also tested the performance of Normalized Google Distance (NGD) and Similarity Ratio (Simratio), which had previously been reported to be the best performing algorithms in an alignment-free remote homology detection benchmark[2]. Between $k$=1 (amino acid composition) and $k$=10, SHARK-score ($T$) consistently offered the best overall performance according to the area under the receiver operating characteristic. By tuning the similarity threshold $T$, SHARK-scores were able to achieve high performance at higher $k$'s where existing metrics begin to deteriorate. A small benchmark notwithstanding, this indicated the promise of SHARK-scores in assessing homology between IDRs. For each algorithm, only $k$-mer lengths where the auPRC would not be over-estimated due to the lack of similar/identical $k$-mers between homologs were plotted (see Methods).

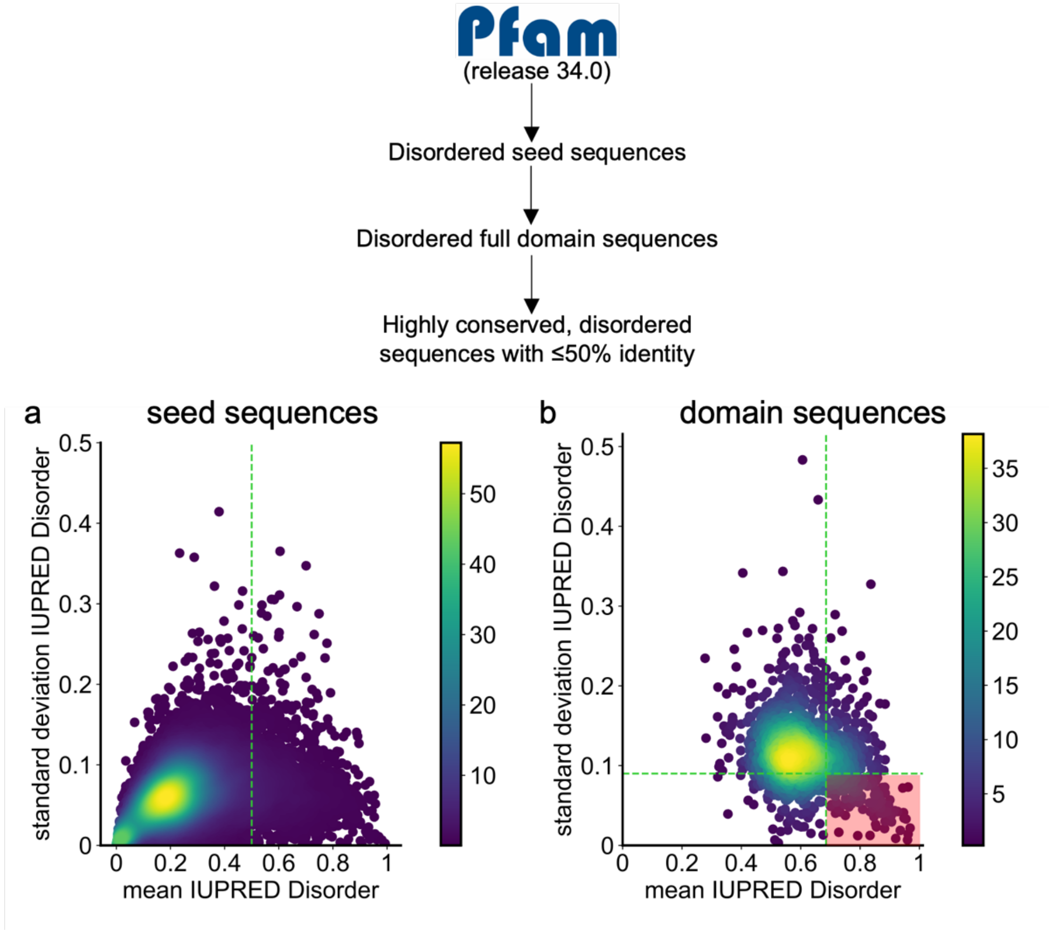


**Figure S3. Overview of the curation of the most disordered Pfam families (alignable-disorder dataset)**. a. Density plot of the mean and standard deviation of seed family disorder of Pfam family seed sequences in the 18188 families analyzed . b. Density plot of the mean and standard deviation of full family disorder of Pfam family domain sequences at max 50% identity. Shaded in red is an area of highly conserved disorder, and contains only families that are in the 75$^{th}$ percentile of mean family disorder and within the 25$^{th}$ percentile of standard deviation, to ensure that all sequences of the family are consistently disordered. This corresponds to 143 families that were included in the alignable-disorder dataset.

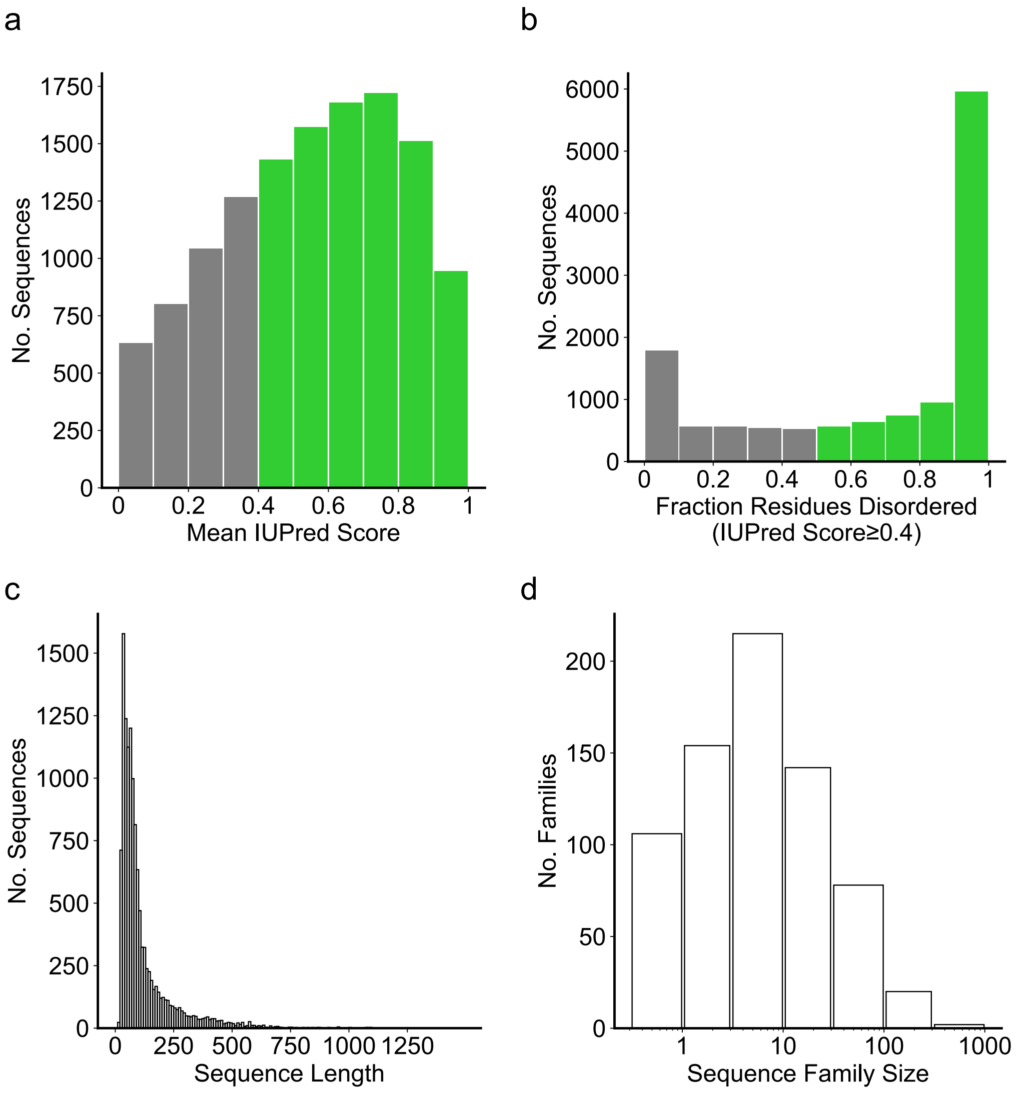


**Figure S4. The unalignable orthologs dataset is enriched in sequences with disordered regions of varying lengths and sequence family sizes**. a. Mean disorder of each sequence in the dataset, with a mean of 0.55 and median of 0.57, despite having no explicit disorder requirement for these non-domain, unalignable sequence. 8857/12602 (70.3%) sequences are considered disordered on average. b. Similarly, 8769/12602 (69.6%) of sequences contain extensive regions of disorder where >50% of residues are predicted to be disordered. c. Sequences are of varying lengths ranging from 15 to 1466 residues long with a mean length of 119. d. Sequence families are of various sizes after ≤50% identity filtering.

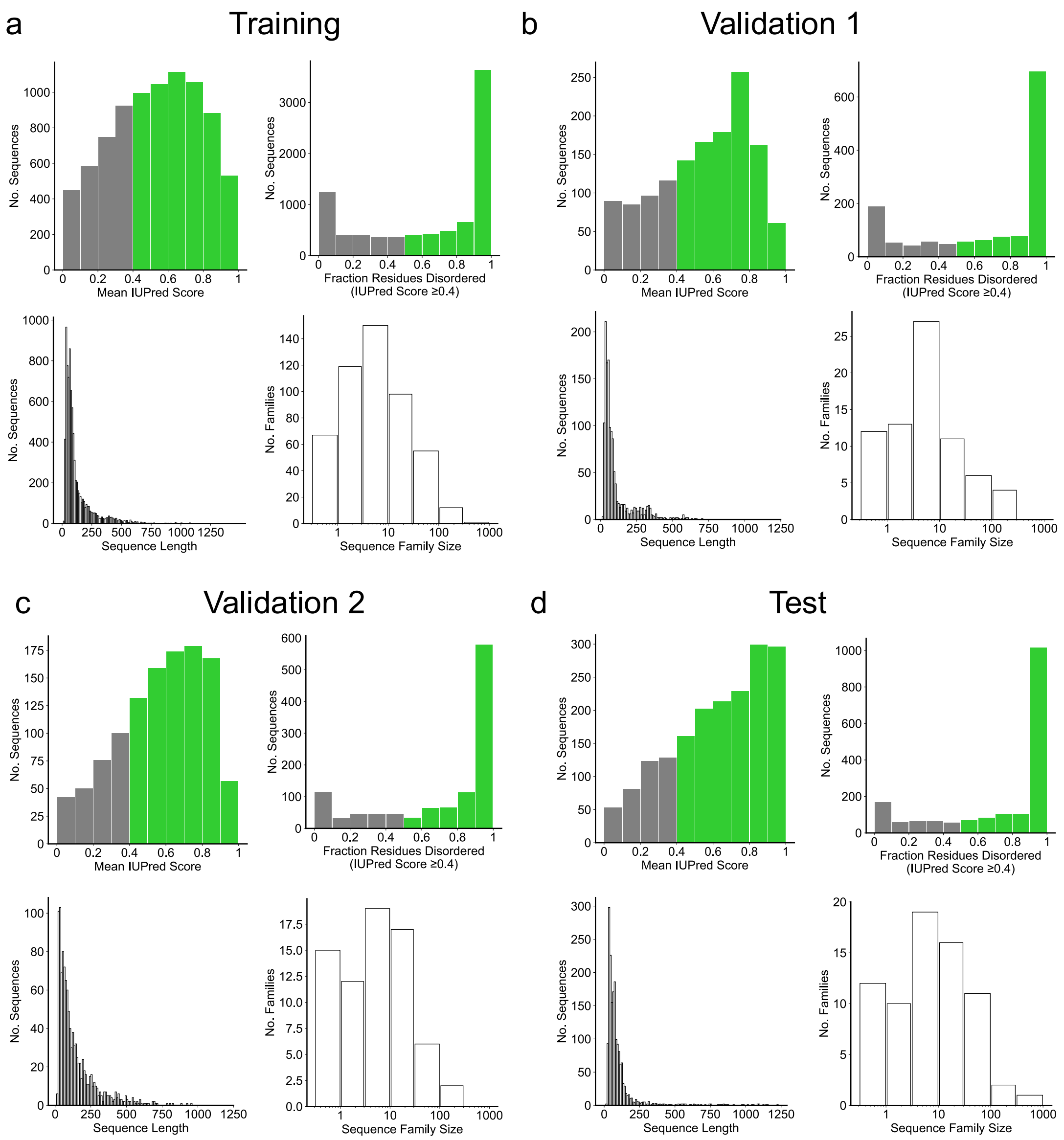


e

| Dataset | Sequences | Sequence Families | True Homolog Pairs (TP) |
|---|---|---|---|
| Training | 8327 | 502 | 709065 |
| Validation 1 | 1353 | 73 | 131031 |
| Validation 2 | 1137 | 71 | 92947 |
| Test | 1785 | 71 | 322581 |

**Figure S5. Disordered sequences are enriched in the unalignable-ortholog sequence datasets**. For each sequence subset- training (a), validation1 (b), validation2 (c) and test (d):

Top left: Distribution of mean sequence disorder, which contains 68%, 71%, 76% and 78% sequences with mean disorder ≥0.4 respectively (shown in green).

Top right: Each dataset contains sequences with extensive regions of disorder where >50% of residues are predicted to be disordered (shown in green).

Bottom right: Each dataset contains sequence families of various sizes.

Bottom left: Sequence length variation is represented across all Disprot sequence subsets.

e. Summary statistics of each unalignable-ortholog sequence dataset.

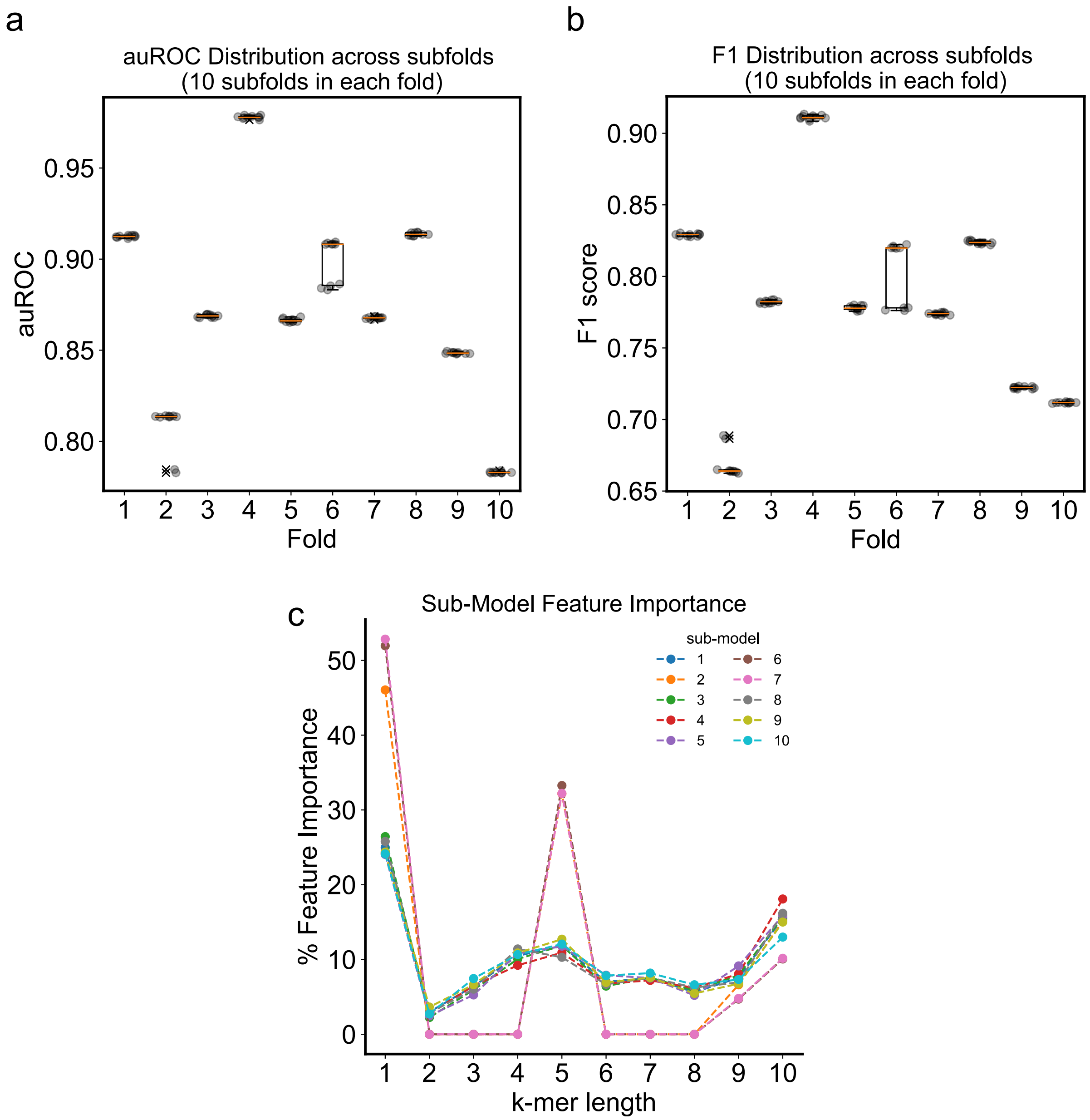


**Figure S6. Performance and feature importance are consistent across sub-folds and sub-models.** Since the number of unrelated sequences (true negatives, TN) far exceeds the number of true homologs (true positives, TP), we trained 10 sub-folds for each sub-model. Each sub-fold contains a training and validation set: the set of true homologs are identical across the sub-folds but the set of unrelated sequences is different. To assess the impact of different unrelated sequences on model training, the area under the receiver operating characteristic curve (auROC, a) and F1 score performance (b) of each sub-fold model on its respective validation dataset was calculated. Since the performance is largely similar, we concluded that the choice of unrelated sequence pairs does not matter significantly (<0.05 difference F1/auROC between min and max). Nonetheless, for each fold we chose the best sub-fold on the validation1 early-stopping dataset used to prevent over-fitting, this is known as a sub-model (i.e. 10 sub-models altogether form the SHARK-dive model). Boxplots show median (orange line) and quartiles, outliers are shown as 'x'. Individual auROC and F1 metrics are shown as grey circles c. Feature importance of each sub-model shows similar trends, with $k$=1, 5 and 10 being the most important in all sub-models.

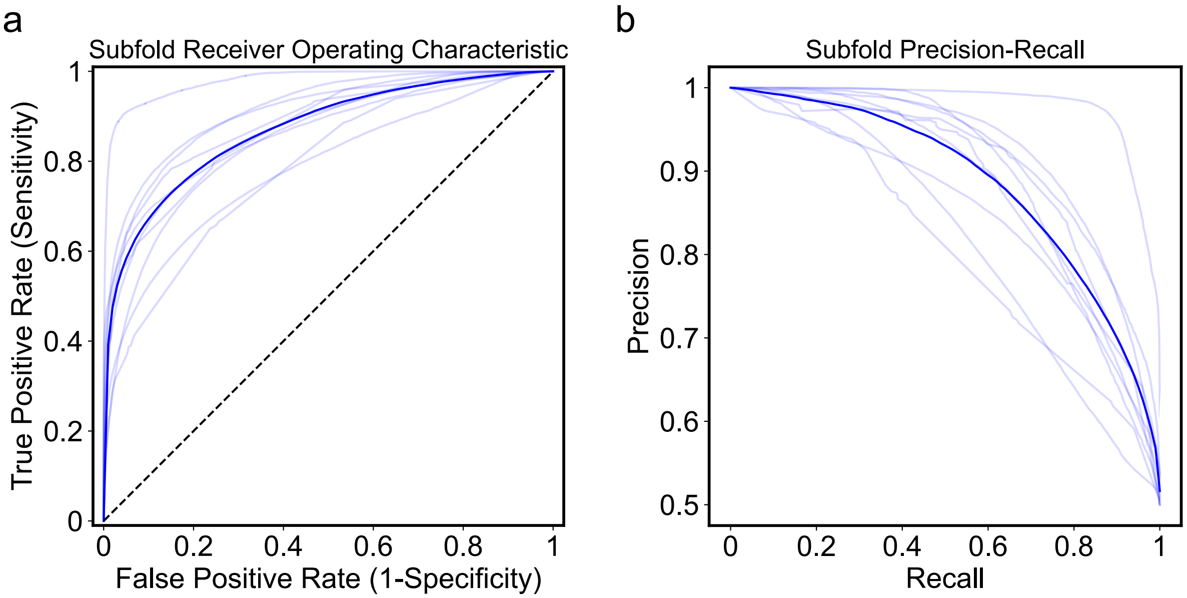


**Figure S7. Ensemble sub-models show consistent performance on validation datasets.** Receiver Operating Characteristic (a) and Precision-Recall (b) curves for each sub-model on its validation dataset. Each ensemble sub-model has a specific validation dataset with withheld sequences not used during model training to assess the performance of the model. Despite variations in sub-model performance, they all show superior performance to a random classifier (dotted line, ROC plot). Differences are due to differences in number of sequences used because sequence family sizes are different, but each dataset is nonetheless balanced in the number of true homologs (TP) and unrelated sequences (TN). There are between 451-453 training families and 49-51 validation families in each fold.

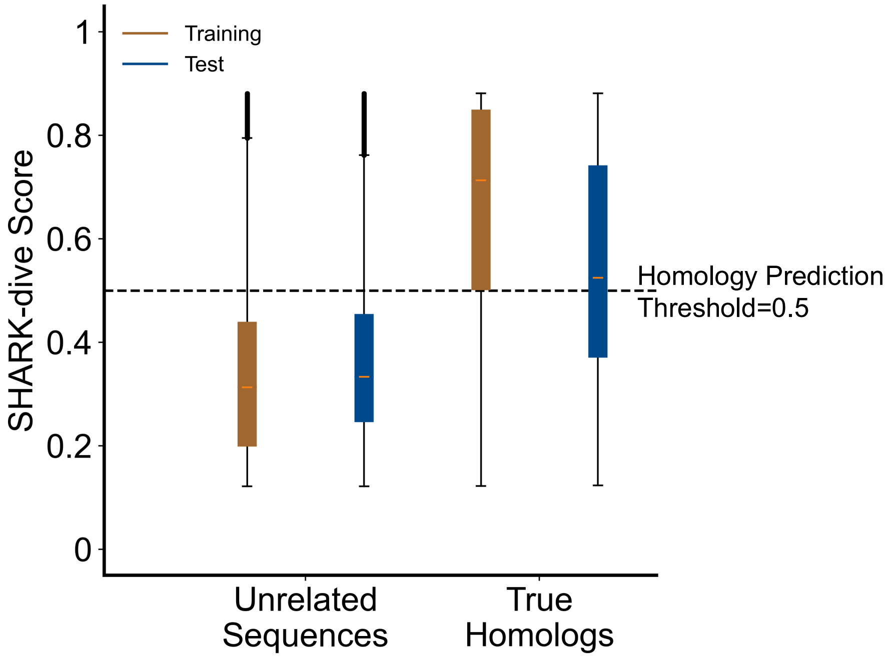


**Figure S8. SHARK-dive distinguishes between true homologs and unrelated sequences in unalignable ortholog sequences.** Boxplot distribution of SHARK-scores on the unalignable orthologs training and test sequences. Boxplots show median (orange line) and quartiles, outliers are shown as ‘x’. The discriminatory power on the withheld test dataset indicates the model had not been overfitted. Orange lines in the boxes indicate the median, where in both datasets the median of unrelated sequences is below the threshold (0.5) whereas the median of the true homologs lies above the threshold.

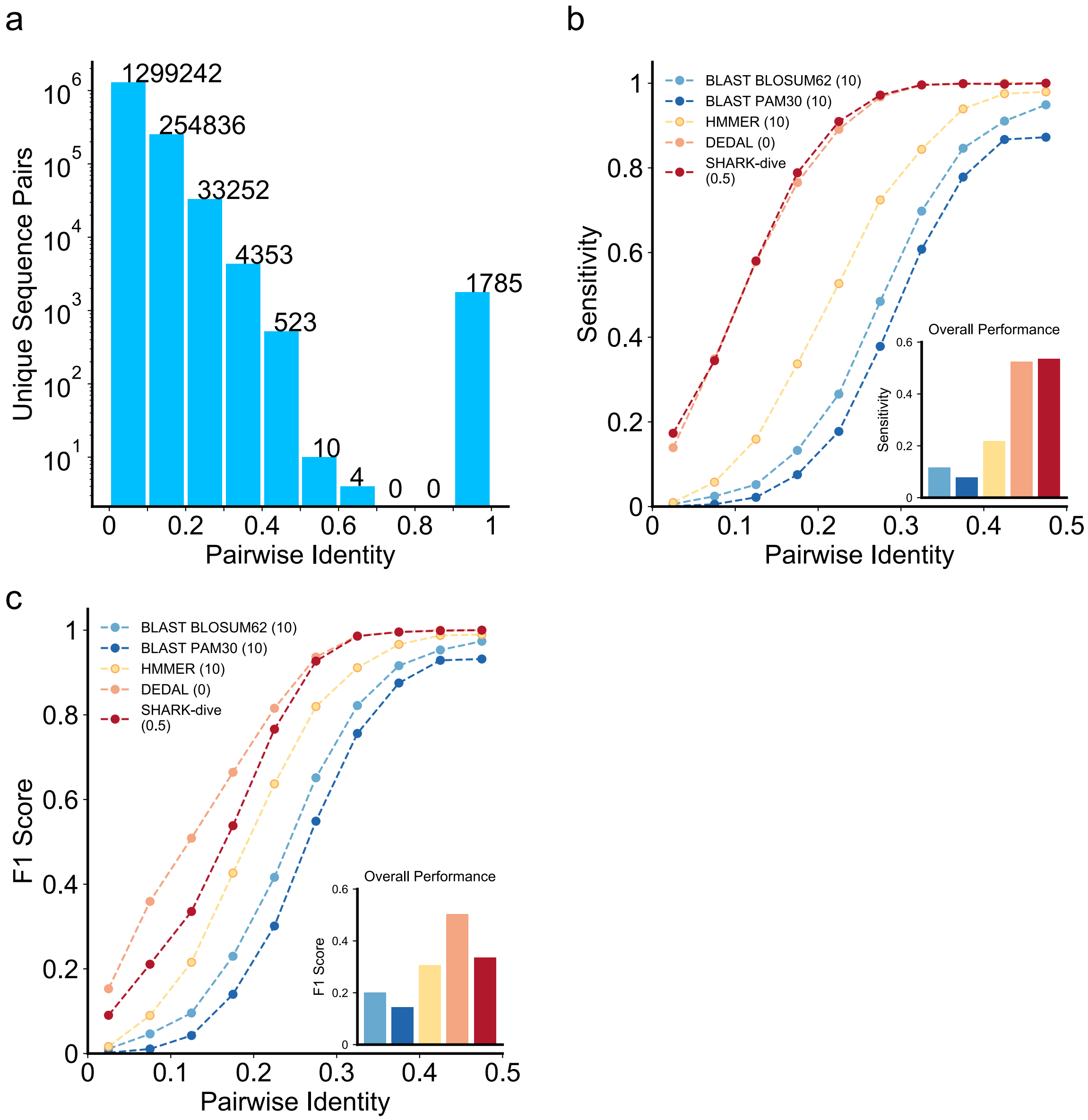


**Figure S9. SHARK-dive achieves high sensitivity to remote, unalignable homologs.** a. The unalignable-ortholog test sequence pairs share low identity. Pairwise identity (PID) in the unalignable orthologs test set after CD-HIT filtering. PID calculated as number of identical amino acids in alignment divided by alignment length (performed using Needleman-Wunsch with BLOSUM62 matrix with default gap penalties (see Methods). Sensitivity (b) and F1-score (c) performance of SHARK-dive and other homology assessment tools on the test dataset, stratified by PID, with the homology detection threshold shown in brackets. SHARK-dive achieves highest sensitivity across most PID bins, with DEDAL offering similar performance. Conventional alignment-based tools such as BLAST and HMMER perform poorly in detecting remote unalignable homologs, even with a highly lenient E-value threshold of 10.

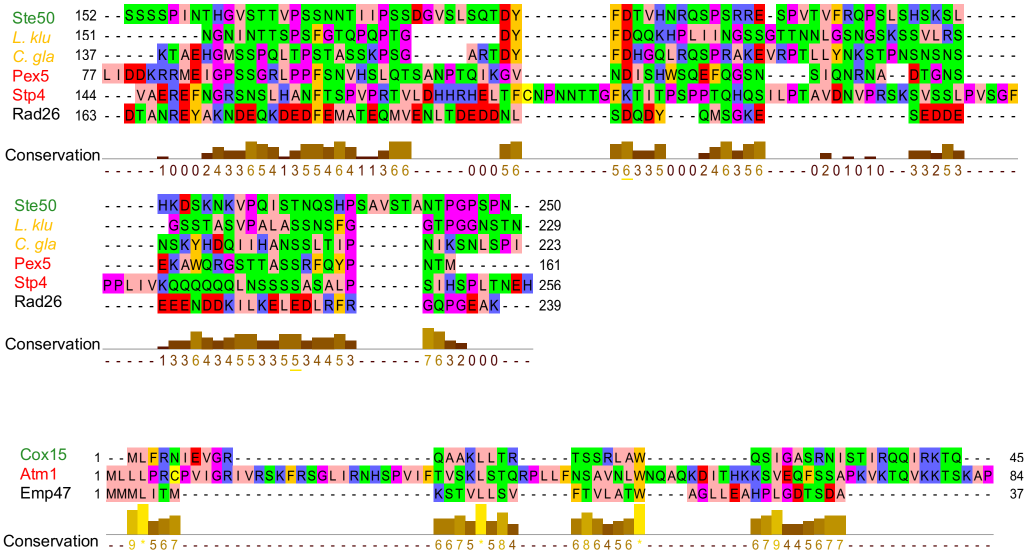


**Figure S10. IDRs with low sequence conservation and poor alignment quality can be functionally homologous.** Multiple Sequence Alignments of Ste50 and Cox15 with their replaced IDRs show poor alignment as indicated by the number of gaps, and the overall low conservation value (as reported in JalView).

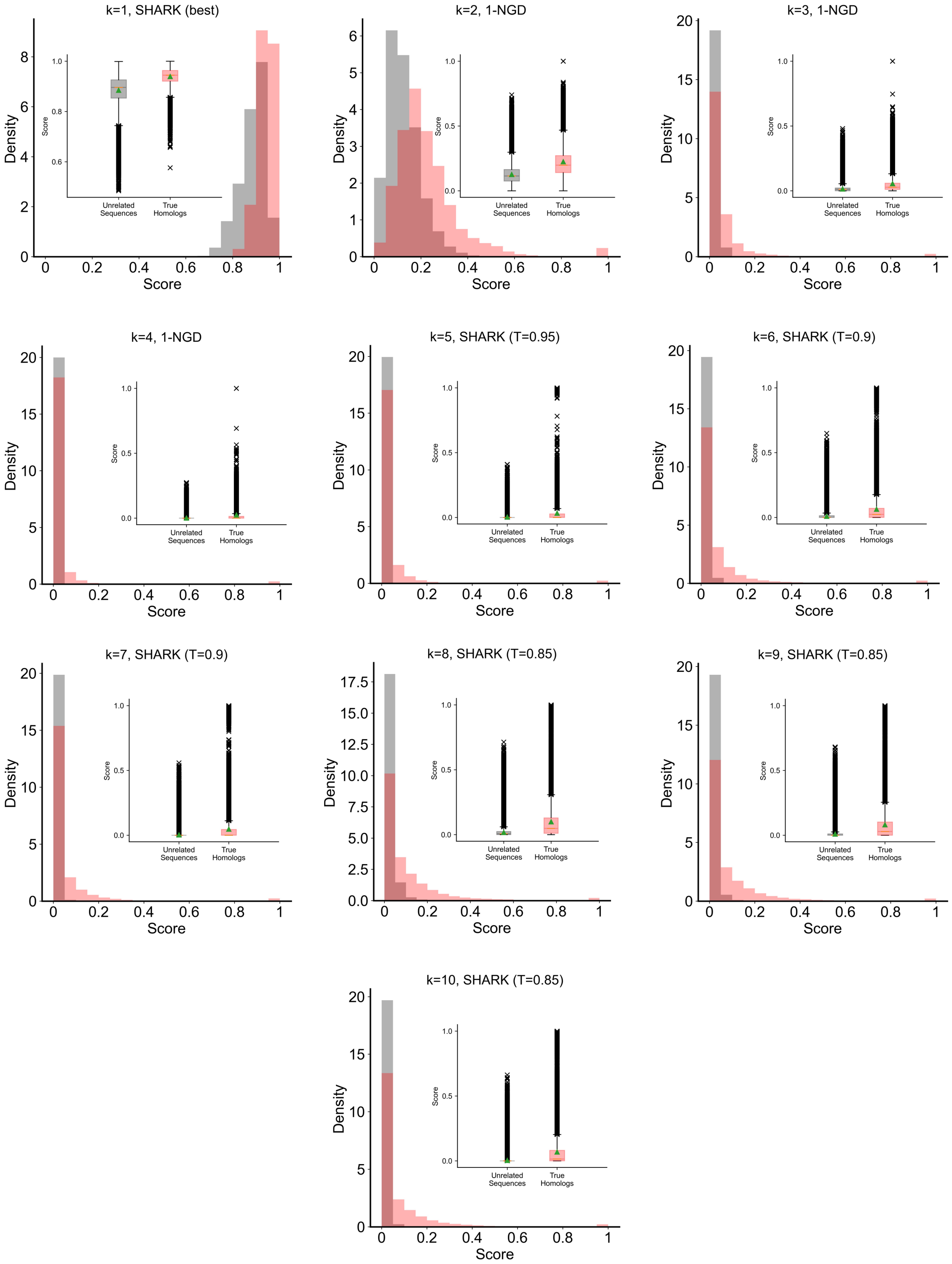


**Figure S11. Individual SHARK-dive features provide discriminatory power over true homologs and unrelated sequences**. Histogram distributions in the unalignable orthologs training set between true homologs and unrelated sequences, with boxplot distributions shown in inset (showing median (orange line), mean (green triangle) and quartiles, outliers as ‘x’). Note: for *k*=2-4, NGD scores are visualized as 1-NGD (otherwise known as the Normalized Google Similarity) such that higher scores represent higher similarity for ease of interpretation.

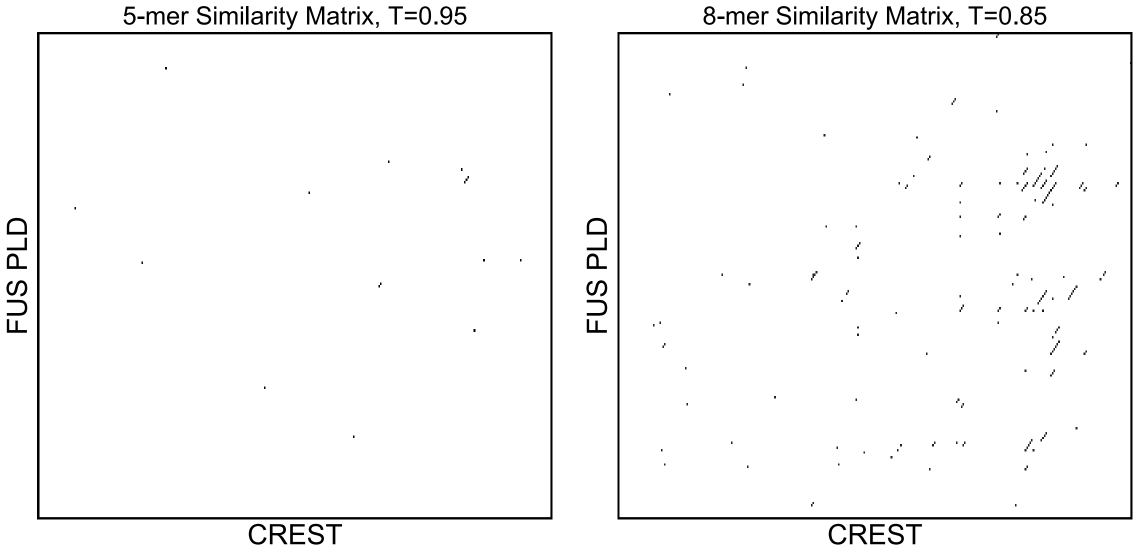


**Figure S12. FUS PLD shares multiple similar *k*-mers with CREST IDR.** Dot plot of similar *k*-mers ($k$=5, $T$=0.95; $k$=8, $T$=0.85) between FUS PLD and CREST (shown as black tiles). These can be mapped back onto each sequence their correspondence visualized (Fig. 5e) to identify regions of similarity between both sequences. Since the decomposition into *k*-mers removes collinear constraints, similar regions across different parts of the IDRs in different positions can be identified.

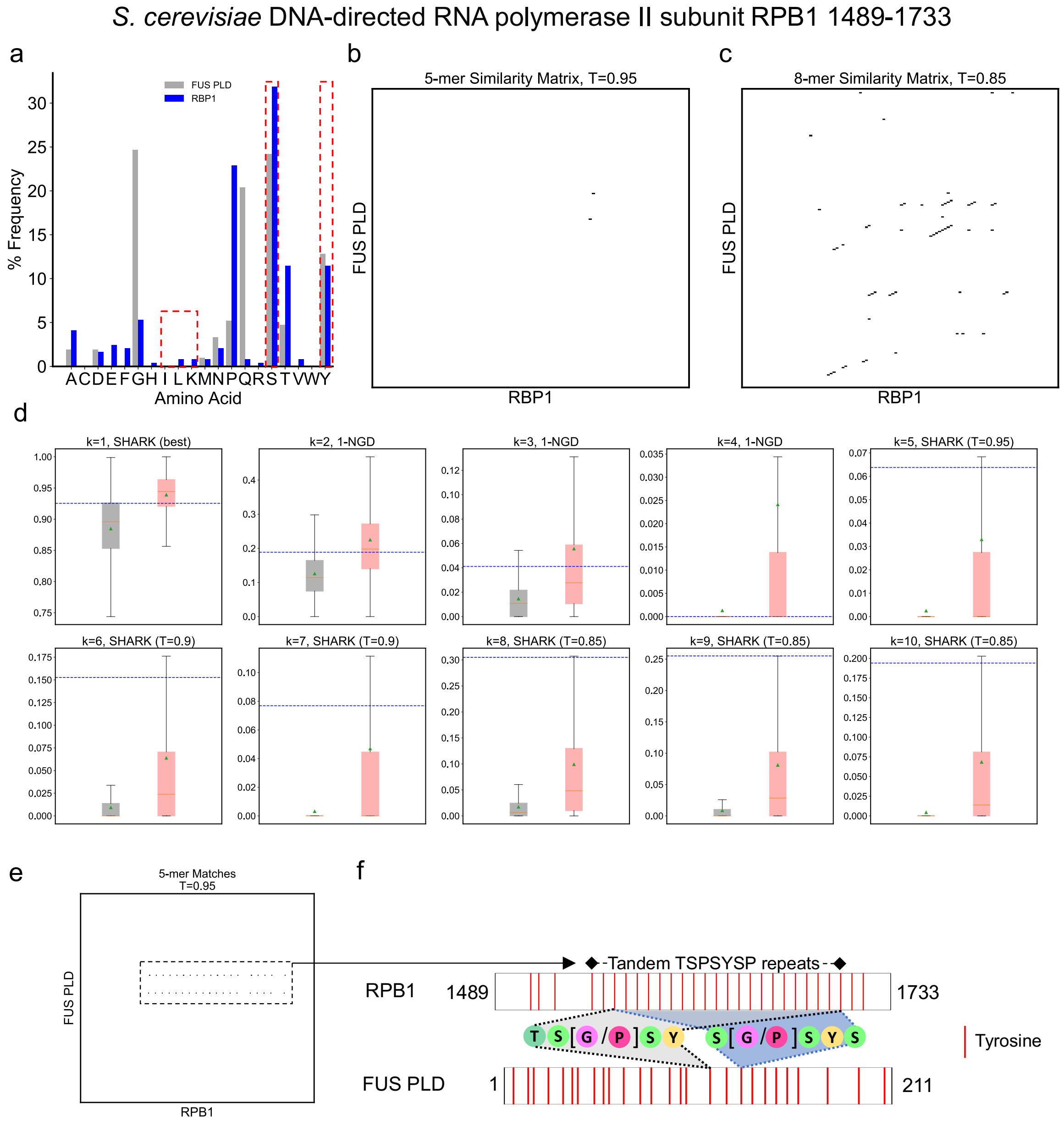


**Figure S13. SHARK-dive predicts a yeast IDR to be homologous to human FUS PLD.** SHARK-dive can also be used to predict homologous IDRs in other species, such as the RPB1 IDR in *S. cerevisiae*, where there are no known orthologs but various pathways where FUS is involved are shared between the two species[3,4]. Compared to the CREST IDR (see Fig. 5c,d), RPB1 shares lower compositional similarities to FUS PLD, as reflected by the lower compositional (*k*=1) score (d). Nonetheless, it does share multiple similar *k*-mers to FUS PLD (*k*=5 and *k*=8 shown in panels b and c respectively). Notably, mapping the highly similar 5-mers (black dots in b) back onto their respective sequences (e), reveals that the 5-mer matches along the sequences map onto a highly repetitive and functional region in RPB1 (f) which would be difficult to align, which contributes to its higher 5-mer score. Whilst only *k=5,* T=0.95 and *k*=8, *T*=0.85 similarity matrices are shown, others can be generated by using the Jupyter Notebook (git.mpi-cbg.de/tothpetroczylab/shark/notebooks/.

Note: In panel d, for *k*=2-4, NGD scores are visualized as 1-NGD (otherwise known as the Normalized Google Similarity) such that higher scores represent higher similarity for ease of interpretation. Boxplots show median (orange line), mean (green triangle) and quartiles, outliers are shown as ‘x’.

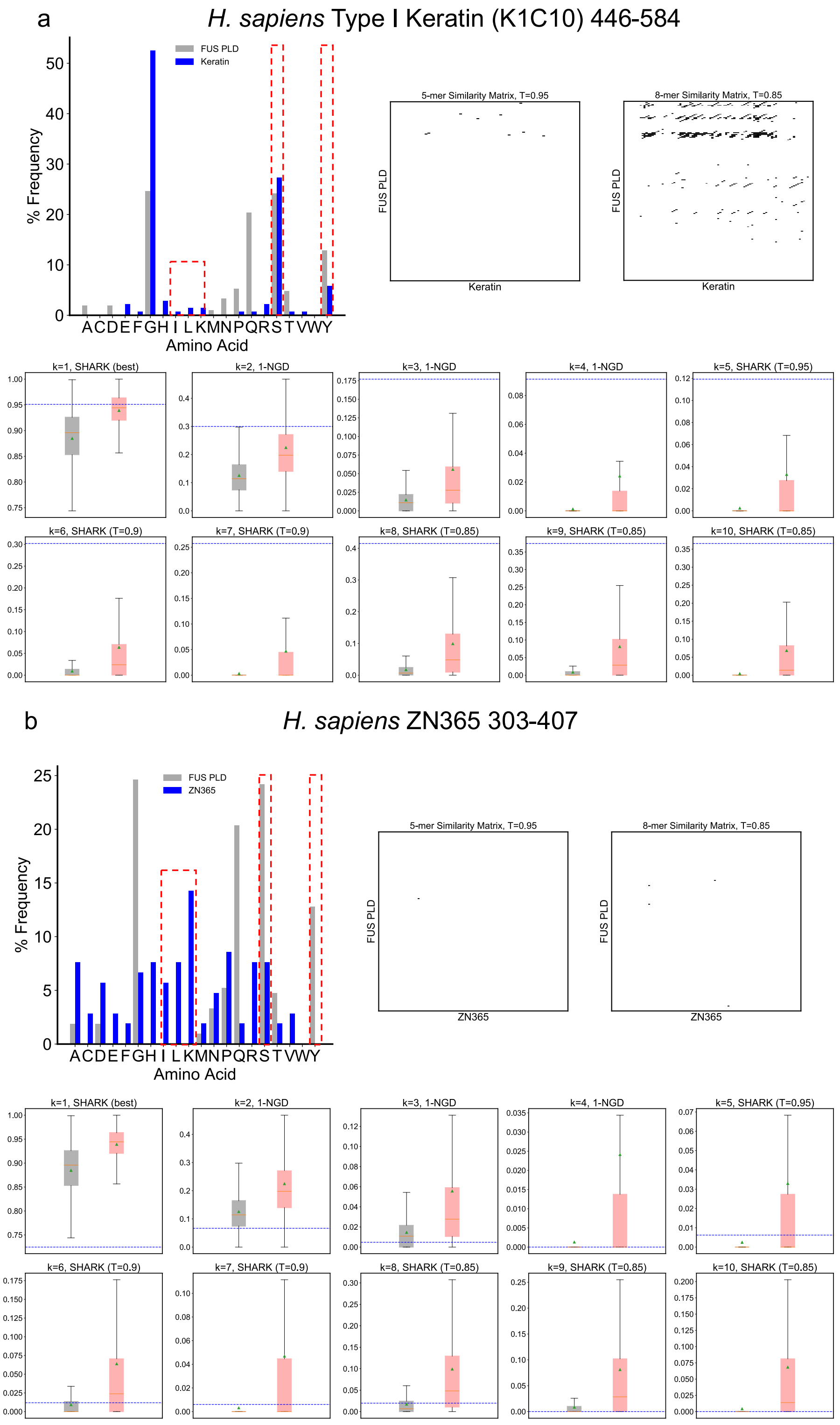


**Figure S14. Predicted homologs and predicted unrelated sequences are highly dissimilar in sequence properties.** FUS PLD (1-211) is searched against a database of IDRs across various proteomes. Shown here is an example of a predicted homolog (a C-terminal IDR of K1C10, a), as well as a predicted non-homolog (ZN365 IDR, b). For each IDR the amino acid composition is plotted against that of FUS PLD. K1C10 shows similar amino acid compositions to FUS, particularly in terms of tyrosine, serine abundance as well as depletion of aliphatics such as leucine and isoleucine, and a lack of lysines. On the contrary, ZN365 has high abundance of K, I and L whilst containing fewer serines and no tyrosine. A similar trend is reflected in the similarity matrices where there are more highly similar, matched sequences to FUS PLD for the predicted homologs than ZN365. Altogether, this is reflected by the difference in *k*-mer scores between the two sequences (see boxplots), where K1C10 *k*-mer scores are higher than most predicted homologs whilst ZN365 *k*-mer scores are closer to the distribution for unrelated sequences. Accordingly, K1C10 is predicted to be homologous to the FUS PLD whereas ZN365 is not. For *k*=2-4, NGD scores are visualized as 1-NGD (otherwise known as the Normalized Google Similarity) such that higher scores represent higher similarity for ease of interpretation. Boxplots show median (orange line), mean (green triangle) and quartiles. Outliers are not shown but can be found in Fig. S11.

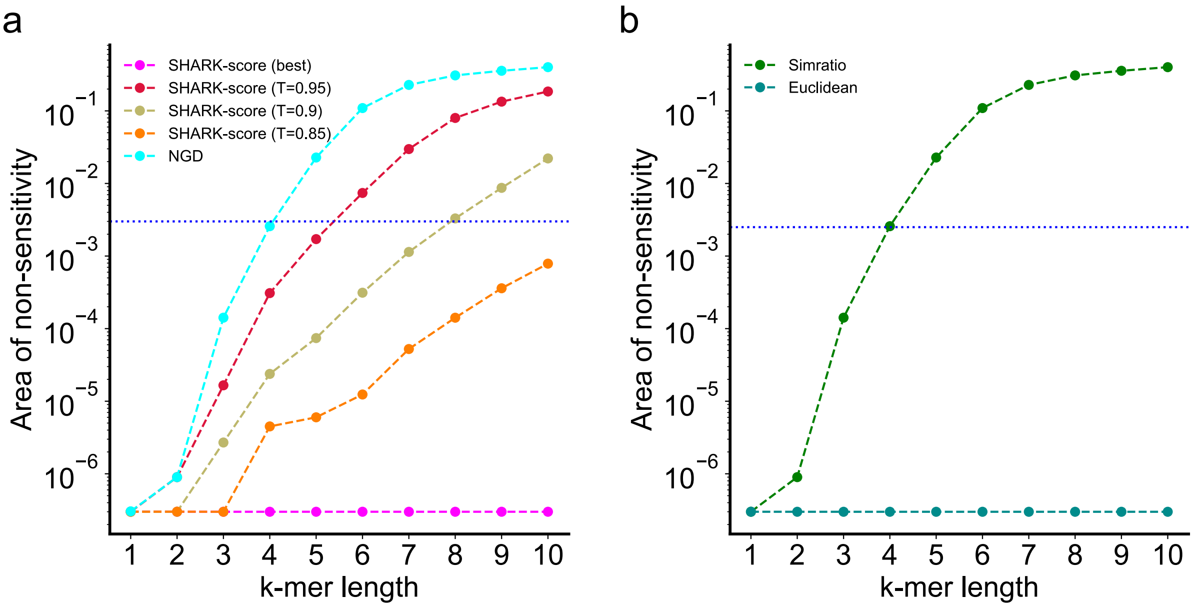


c

| Alignment-free algorithm | Maximum *k*-mer length |
|---|---|
| SHARK-score (best) | 10 |
| Normalized Google Distance (NGD) | 4 |
| SHARK-score (T=0.95) | 5 |
| SHARK-score (T=0.9) | 7 |
| SHARK-score (T=0.85) | 10 |
| Similarity Ratio (Simratio) | 4 |
| Euclidean | 10 |

**Figure S15. Area of non-sensitivity in precision-recall curve for the alignable-disorder dataset.** Area calculated via trapezium rule between the final recall point (recall=1.0) and the previous recall point for various alignment-free algorithms (a,b). This is to identify the range in which the scores are unable to distinguish between sequences in the same PFAM family (true homologs) and unrelated sequences because the maximum distance value is reached. Sequence comparison algorithms at *k*-mer lengths that have non-sensitivity area >0.003 (threshold, dotted blue line) are not considered in feature selection nor have auPRC calculated since the auPRC would be overestimated, the maximum *k*-mer length considered is summarized in c).

### Supplementary Tables

**Table S1. Parameters used for benchmarking Smith-Waterman local alignment**

| Substitution Matrix | Gap Opening Penalty (*G*) [5] | Gap Extension Penalty (*L*) | Notes | Ref |
|---|---|---|---|---|
| BLOSUM62 | -11 | -1 | BLASTp default parameters | 6–8 |
| PAM30 | -9 | -1 | BLASTp-short default parameters | |
| PFASUM70 | -15 | -1.5 | generated using DECIPHER package in R and rounded to nearest integer as per Keul *et al*.[9] | 9 |
| EDSSMat60 | -6 | -2 | Medium Disorder (MD) | 10 |
| EDSSMat80 | -5 | -2 | | |
| EDSSMat50 | -6 | -2 | | |
| EDSSMat90 | -5 | -2 | | |
| EDSSMat62 | -5 | -2 | | |
| EDSSMat70 | -5 | -2 | | |
| EDSSMat75 | -5 | -2 | | |
| EDSSMat60 | -14 | -3 | High Disorder (HD) | |
| EDSSMat80 | -15 | -3 | | |
| EDSSMat50 | -18 | -2 | | |
| EDSSMat90 | -19 | -2 | | |
| EDSSMat62 | -19 | -2 | | |
| EDSSMat70 | -19 | -2 | | |
| EDSSMat75 | -19 | -2 | | |

**Note:** Matrices shaded in grey were also used in benchmarking the alignable-disorder dataset

**Table S2. The best performing alignment-free algorithm selected for each *k*-mer length.**

| *k* | Algorithm |
|---|---|
| 1 | SHARK-score (best) |
| 2 | NGD |
| 3 | NGD |
| 4 | NGD |
| 5 | SHARK-score (T=0.95) |
| 6 | SHARK-score (T=0.9) |
| 7 | SHARK-score (T=0.9) |
| 8 | SHARK-score (T=0.85) |
| 9 | SHARK-score (T=0.85) |
| 10 | SHARK-score (T=0.85) |

**Note:** NGD: Normalized Google Distance.

**Table S3. Benchmarking against Local Alignment (Smith-Waterman algorithm) on alignable-disorder dataset**

| Method, Threshold | Recall | Specificity | Precision | Accuracy | f1 | auPRC |
|---|---|---|---|---|---|---|
| SHARK-dive, 0.5 | **0.536** | 0.814 | 0.245 | 0.786 | **0.336** | **0.344** |
| EDSSMat60 (MD), 35 | 0.198 | 0.956 | 0.335 | 0.879 | 0.249 | 0.258 |
| EDSSMat80 (MD), 37 | 0.200 | 0.954 | 0.332 | 0.878 | 0.250 | 0.256 |
| EDSSMat50 (MD), 36 | 0.197 | 0.956 | 0.336 | 0.879 | 0.249 | 0.256 |
| EDSSMat90 (MD), 37 | 0.198 | 0.956 | 0.337 | 0.879 | 0.250 | 0.256 |
| EDSSMat62 (MD), 37 | 0.202 | 0.953 | 0.326 | 0.877 | 0.250 | 0.256 |
| EDSSMat70 (MD), 37 | 0.201 | 0.954 | 0.328 | 0.877 | 0.249 | 0.256 |
| EDSSMat75 (MD), 37 | 0.202 | 0.953 | 0.325 | 0.877 | 0.249 | 0.256 |
| EDSSMat60 (HD), 34 | 0.170 | 0.973 | 0.417 | 0.892 | 0.242 | 0.253 |
| EDSSMat80 (HD), 34 | 0.173 | 0.972 | 0.407 | 0.891 | 0.242 | 0.253 |
| EDSSMat62 (HD), 34 | 0.173 | 0.971 | 0.402 | 0.890 | 0.242 | 0.253 |
| EDSSMat70 (HD), 34 | 0.172 | 0.971 | 0.404 | 0.890 | 0.242 | 0.253 |
| EDSSMat75 (HD), 34 | 0.173 | 0.971 | 0.400 | 0.890 | 0.241 | 0.253 |
| EDSSMat90 (HD), 34 | 0.171 | 0.973 | 0.412 | 0.891 | 0.241 | 0.252 |
| EDSSMat50 (HD), 34 | 0.175 | 0.969 | 0.388 | 0.889 | 0.241 | 0.252 |
| PAM30, 35 | 0.244 | 0.916 | 0.246 | 0.848 | 0.245 | 0.250 |
| PFASUM70, 41.5 | 0.376 | 0.792 | 0.169 | 0.750 | 0.233 | 0.238 |
| BLOSUM62, 31 | 0.341 | 0.819 | 0.175 | 0.771 | 0.231 | 0.237 |

**Note:** For completeness, specificity and accuracy values are included although we note that these metrics, much like the auROC, are not suitable for our imbalanced test dataset due to the high number of unrelated pairs from an all-vs-all comparison. Table is sorted in descending order of auPRC, with highest auPRC and F1 shown in bold. Shaded in blue are the precision-recall curves shown in Fig. 3d.

**Table S4. Tools and parameters used for benchmarking against alignment-based homology search tools**

| Tool | Substitution Matrix | Gap Opening Penalty (*G*) [5] | Gap Extension Penalty (*L*) | Thresholds tested | Notes | Ref. |
|---|---|---|---|---|---|---|
| BLASTp | BLOSUM62 | -11 | -1 | E-value: 10,1,0.05 | BLASTp default parameters | [6,11] |
| | PAM30 | -9 | -1 | | | |
| | PAM30 | -9 | -1 | | BLASTp-short (no composition-based statistics, word_size=2) | |
| | BLOSUM50 | -13 | -2 | | | |
| | PAM250 | -14 | -2 | | | |
| | BLOSUM90 | -10 | -1 | | | |
| | BLOSUM45 | -15 | -2 | | | |
| | BLOSUM80 | -10 | -1 | | | |
| | PAM70 | -10 | -1 | | | |
| pHMMER | Default | Default | Default | E-value: 10,1,0.001 Bitscore: 7 | --max option used for maximum sensitivity | [12,13] |
| DEDAL (homology classifier) | - | - | - | 0 | DEDAL homology classifier logit threshold | [14] |

**Note:** For BLASTp and pHMMER reporting E-values were set arbitrarily high ($10^4$ for BLASTp, $10^{14}$ for pHMMER)

**Table S5. Benchmarking on the unalignable-orthologs test dataset**

| Method, Threshold | Recall | Specificity | Precision | Accuracy | F1 |
|---|---|---|---|---|---|
| Dedal Logit, 0 | 0.524 | 0.938 | 0.487 | 0.896 | **0.505** |
| SHARK-dive, 0.5 | **0.536** | 0.814 | 0.245 | 0.786 | 0.336 |
| pHMMER E-value, 10 (default reporting E-value) | 0.219 | 0.977 | 0.515 | 0.900 | 0.307 |
| BLAST BLOSUM50, 10 | 0.134 | 0.993 | 0.694 | 0.906 | 0.225 |
| BLAST PAM250, 10 | 0.129 | 0.996 | 0.765 | 0.908 | 0.221 |
| pHMMER bitscore, 7 (default reporting bitscore) | 0.121 | 0.998 | 0.867 | 0.909 | 0.213 |
| pHMMER E-value, 1 (default reporting E-value on webserver) | 0.119 | 0.998 | 0.856 | 0.909 | 0.210 |
| BLAST BLOSUM90, 10 | 0.116 | 0.997 | 0.829 | 0.908 | 0.203 |
| BLAST BLOSUM62, 10 | 0.115 | 0.996 | 0.787 | 0.907 | 0.201 |
| BLAST BLOSUM45 10 | 0.111 | 0.996 | 0.740 | 0.906 | 0.193 |
| BLAST PAM250, 1 | 0.106 | 0.999 | 0.954 | 0.909 | 0.191 |
| BLAST BLOSUM50, 1 | 0.105 | 0.999 | 0.944 | 0.909 | 0.190 |
| BLAST BLOSUM80, 10 | 0.103 | 0.998 | 0.860 | 0.907 | 0.184 |
| BLAST short, 10 | 0.102 | 0.998 | 0.867 | 0.908 | 0.183 |
| BLAST BLOSUM62, 1 | 0.098 | 1.000 | 0.964 | 0.908 | 0.178 |
| BLAST BLOSUM45, 1 | 0.098 | 0.999 | 0.942 | 0.908 | 0.177 |
| BLAST BLOSUM90, 1 | 0.095 | 1.000 | 0.974 | 0.908 | 0.173 |
| BLAST PAM70, 10 | 0.094 | 0.999 | 0.917 | 0.907 | 0.171 |
| BLAST BLOSUM80, 1 | 0.089 | 1.000 | 0.978 | 0.908 | 0.164 |
| BLAST short, 1 | 0.084 | 1.000 | 0.980 | 0.907 | 0.155 |
| BLAST PAM70, 1 | 0.083 | 1.000 | 0.988 | 0.907 | 0.153 |
| BLAST BLOSUM45, 0.05 | 0.081 | 1.000 | 0.995 | 0.907 | 0.151 |
| BLAST PAM250, 0.05 | 0.081 | 1.000 | 0.997 | 0.907 | 0.150 |
| BLAST BLOSUM50, 0.05 | 0.081 | 1.000 | 0.996 | 0.907 | 0.149 |
| BLAST BLOSUM62, 0.05 | 0.078 | 1.000 | 0.997 | 0.907 | 0.145 |
| BLAST PAM30, 10 | 0.078 | 0.999 | 0.938 | 0.906 | 0.145 |
| BLAST BLOSUM90, 0.05 | 0.074 | 1.000 | 0.998 | 0.906 | 0.138 |
| BLAST BLOSUM80, 0.05 | 0.072 | 1.000 | 0.998 | 0.906 | 0.135 |
| BLAST PAM30, 1 | 0.071 | 1.000 | 0.989 | 0.906 | 0.132 |
| BLAST PAM70, 0.05 | 0.068 | 1.000 | 0.999 | 0.906 | 0.127 |
| BLAST short, 0.05 | 0.066 | 1.000 | 0.998 | 0.905 | 0.124 |
| pHMMER E-value, 0.001 | 0.064 | 1.000 | 0.999 | 0.905 | 0.121 |
| BLAST PAM30, 0.05 | 0.060 | 1.000 | 0.999 | 0.905 | 0.113 |

**Note:** For completeness, specificity and accuracy values are included although we note that these metrics, much like the auROC, are not suitable for our imbalanced test dataset due to the high number of unrelated pairs from an all-vs-all comparison. Table is sorted in descending order of F1, with highest recall (sensitivity) and F1 shown in bold. Shaded in blue are thresholds shown in Fig. 3e and f.

**Table S6. Pairwise identity of sequences compared in Fig. 4**

| Query | Entry | Pairwise Identity |
|---|---|---|
| sp\|P25344\|STE50_YEAST/152-250 | sp\|P35056\|PEX5_YEAST/77-161 | 0.094 |
| sp\|P25344\|STE50_YEAST/152-250 | sp\|Q07351\|STP4_YEAST/144-256 | 0.133 |
| sp\|P25344\|STE50_YEAST/152-250 | sp\|P40352\|RAD26_YEAST/163-239 | 0.018 |
| sp\|P25344\|STE50_YEAST/152-250 | STE50_ORTHOLOG/C.gla/137-223 | 0.248 |
| sp\|P25344\|STE50_YEAST/152-250 | STE50_ORTHOLOG/L.klu/151-229 | 0.154 |
| sp\|P40086\|COX15_YEAST/1-45 | sp\|P40416\|ATM1_YEAST/1-84 | 0.111 |
| sp\|P40086\|COX15_YEAST/1-45 | sp\|P43555\|EMP47_YEAST/1-37 | 0.013 |

**Table S7. Reference proteomes obtained from UniProt 2022-05 Release**

| Unique proteome ID | Organism (common name) | Taxonomy ID | No. IDRs |
|---|---|---|---|
| UP000005640 | Homo Sapiens (Human) | 9606 | 69383 |
| UP000000589 | Mus musculus (Mouse) | 10090 | 67570 |
| UP000000625 | Escherichia coli | 83333 | 5207 |
| UP000002311 | Saccharomyces cerevisiae (Baker's yeast) | 559292 | 14985 |
| UP000006548 | Arabidopsis thaliana (Mouse-ear cress) | 3702 | 60185 |
| UP000000437 | Danio rerio (Zebrafish) | 7955 | 89418 |
| UP000000803 | Drosophila melanogaster (Fruit fly) | 7227 | 41643 |

**Table S8. Performance of different tools on synthetic IDRs derived from Cox15 N-terminus**[15]

| WT IDR: | Cox15 | | |
|---|---|---|---|
| Replaced IDR: | N1 | P1 | P2 |
| Experiment (ground-truth) | ✗ | ✓ | ✓ |
| SHARK-dive | ✓ | ✓ | ✓ |
| DEDAL | ✓ | ✓ | ✓ |
| BLAST (BLOSUM62) | ✗ | ✗ | ✗ |
| HMMER | ✓ | ✗ | ✗ |

Note: E-value threshold of <0.05 used for BLAST and HMMER, 0.5 for SHARK-dive, 0 for DEDAL homology classifier. Blue cells indicate concordant homology predictions to experimental results (dark blue).

**Table S9. Version information of all tools and programs**

| Tool | Minimum Version | Other versions used |
|---|---|---|
| Python | 3.6.5 | 3.7.4, 3.7.7 |
| BLAST (makeblastdb, BLASTp) | 2.13.0 | |
| HMMER (pHMMER, HMMscan) | 3.1b2 | |
| MUSCLE | 3.8.31 | |
| Catboost | 1.0.0 | |
| Numpy | 1.15.0 | 1.19.5, 1.21.6 |
| Biopython | 1.71 | 1.75, 1.78, 1.80 |
| Pandas | 1.0.5 | 1.1.2, 1.1.3, 1.3.5 |
| Sci-kit Learn | 0.19.1 | 0.23.1, 0.24.2, 1.0 |
| Scipy | 1.5.4 | 1.6.2, 1.7.1 |
| Matplotlib | 3.1.1 | 3.3.4 |
| Pickle | 4.0 | |
| Alfpy | 1.0.6 | |
| R | 3.6.3 | |
| DECIPHER | 2.14.0 | |
| Tensorflow | 2.12.0-dev20221213 | |
| DEDAL | TF2.0 Saved Model (v3) | |
| CD-HIT | 4.6 | 4.8.1 |

### Supplementary Datasets

**Dataset S1. Alignable-disorder dataset** (Data_S1_alignable_disorder_dataset.zip).

**Dataset S2. Unalignable-ortholog dataset** (Data_S2_unalignable_ortholog_dataset.zip).

**Dataset S3. Homology predictions of FUS PLD (aa 1-211) to IDRs in 7 model organisms** (Data_S3_FUS-PLD_homology_pred.csv)

**Dataset S4. Hyperparameter tuning results** (see Data_S4_hyperparameters.csv)
